# DAWN-SCAPE: Discovering Associations With Networks through Shared Contextual Analysis of Phenotype and Expression

**DOI:** 10.64898/2025.12.09.691744

**Authors:** Maya Shen, Jinjin Tian, Bernie Devlin, Kathryn Roeder

## Abstract

The molecular basis of phenotypes is often explored by contrasting gene expression from relevant tissue taken from individuals classified into phenotypic extremes, such as affected versus unaffected individuals. Analysis of this differential expression (DE) typically identifies many genes of interest. However, it is not clear which genes differ between extremes because they alter phenotypic liability and which show differences as a result of the extreme phenotype itself. We propose a formal model to distinguish between genes that are upstream and “cause” differential expression versus those for which differential expression is a result of the initial manifestation of phenotype. Relying on two sets of *p*-values, one from differential expression analysis and one from gene-specific association with phenotype (AP), and a gene coexpression or other gene-based network that serves as a bridge, our method identifies communities of genes more likely upstream or downstream of the phenotype. Our method consists of three major steps: 1) gene network construction, 2) evaluation of DE and AP signal within the network to infer hidden states, and 3) detection of gene communities. We apply our method to data that were generated to assess the biological basis of autism spectrum disorder (ASD) and Alzheimer’s disease (AD). Our results highlight neuronal and synaptic biology as being upstream of ASD, whereas downstream processes are all non-neuronal. For AD, our results are consistent with existing hypotheses; yet, they also lend support for a recent unifying hypothesis involving cofilin/actin biology.

## Introduction

The molecular basis of phenotypes, especially human diseases and disorders, is often explored by contrasting gene expression profiles of relevant tissue or cells taken from affected and unaffected individuals. For instance, for autism spectrum disorder (ASD), postmortem cortical tissue shows profound differences in expression of thousands of genes, relative to that from neurotypical (NT) individuals, including upregulation of genes involved in inflammatory and immune processes and downregulation of genes involved in synaptic function [1, 2, 3]. Nonetheless, which genes differ because they influence liability for ASD and which differ as a result of ASD is far less clear, and this is true for most if not all studies of differential expression. Informal approaches to distinguish between these two sets of genes usually involve some form of enrichment analysis of genetically associated genes [3]. We propose a more formal model to distinguish between genes that are differentially expressed and affect liability, which we call active genes, from those that are differentially expressed but arise as a result of selection on phenotype, which we will call reactive. We use results from studies of ASD and Alzheimer’s disease (AD) to illustrate the approach, demonstrating its ability to identify groups of genes that are active and reactive, thereby providing insights into the molecular mechanisms underlying the etiology of these two conditions.

Our approach integrates several sources of information involving genes: gene-specific association with phenotype (AP); differential gene expression (DE) or other differential abundance data (e.g., protein); and gene-based networks, such as those derived from gene coexpression profiles or protein-protein interaction. We build on earlier work that sought to expand the set of AP genes by using gene networks under the principle that such genes tend to cluster in neighborhoods of the network [4, 5, 6]. This approach is implemented by jointly modeling a gene network and gene associations using a hidden random Markov field (HMRF) model. To adapt this Discovering Association With Networks (DAWN) model to separate genes that are AP *and* DE (“active” genes) versus those that are solely reactive or likely to be (putative-reactive or “p-reactive” genes), we expand the hidden states of genes in the HMRF model to represent the active and reactive states, as well as another class for genes fitting neither category (other genes). Within appropriately constructed gene networks, the model assumes there are gene communities solely composed of reactive genes, which we label emergent communities, and others composed of active and reactive genes, which we call etiological communities. The new DAWN-SCAPE (Shared Contextual Analysis of Phenotype and Expression) algorithm involves three major steps: 1) gene network construction, 2) evaluation of DE and AP signal within the network using a HMRF model to infer hidden states, and 3) detection of gene communities (Figure 1). We develop novel methods throughout the analytic pipeline to solve methodological and application-specific challenges.

**Figure 1:**
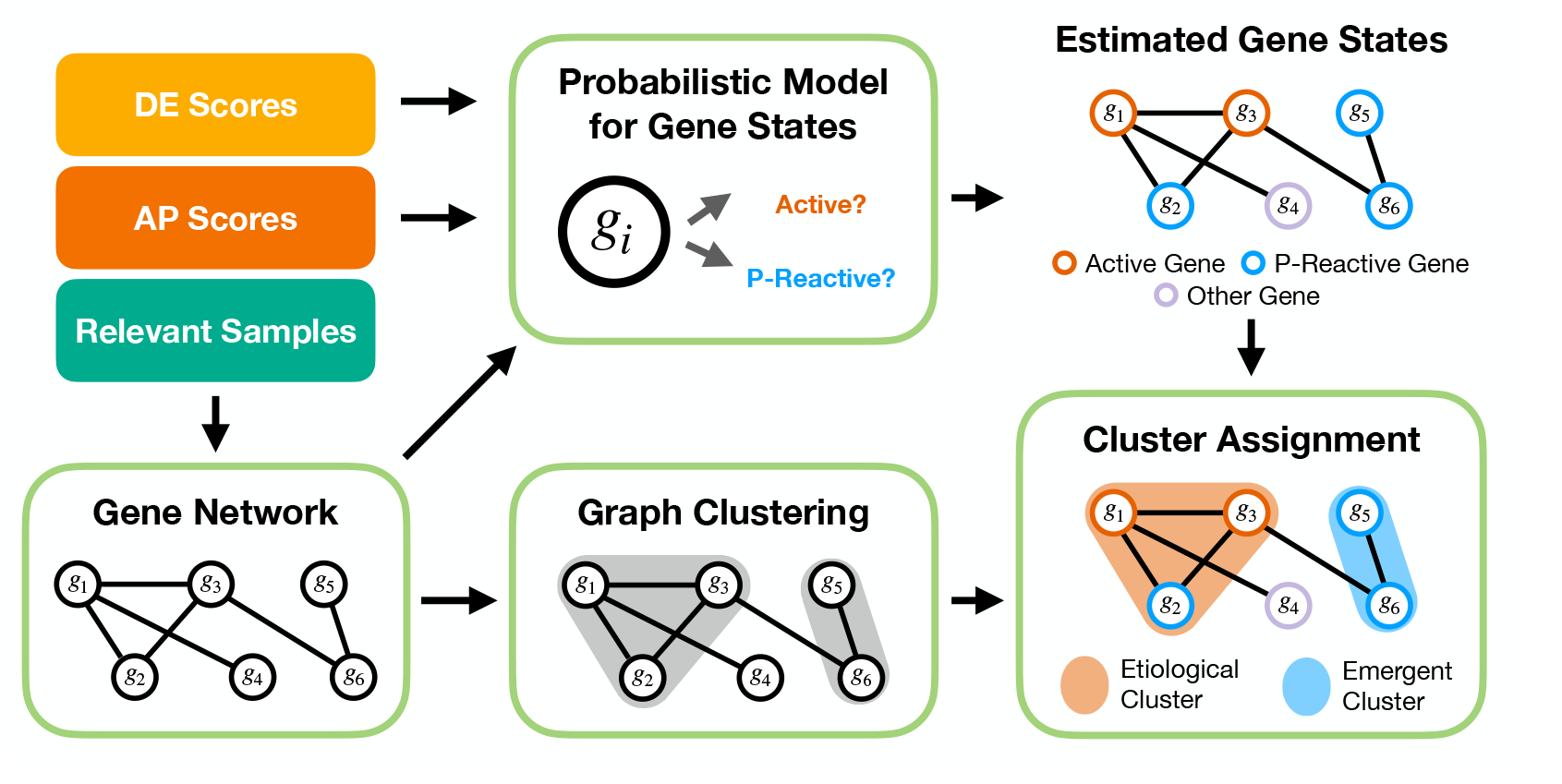
Algorithm to interpret the roles of differentially expressed (DE) genes. Inputs are DE scores, AP scores (e.g., genetic association with phenotype), and relevant information from which to estimate gene network (e.g., gene expression in brain tissue). The final output are genes partitioned into clusters that are defined as etiological and emergent based on the composition of the clusters in terms of the proportion of genes labeled as active (AP *and* DE) versus p-reactive (DE *only*).

## Methods

Methods such as WGCNA [7] assess bivariate dependencies between the expression patterns of all pairs of genes to infer “coexpression networks” and corresponding subnetworks called modules. Such networks are dense in edges between nodes (genes), many of which are not driven by regulatory mechanisms. Alternatively, by identifying independence structure between genes, conditional on all other expression quantities, Gaussian graphical models produce a sparser and more interpretable gene network. Here, we employ one such method, Partial Neighborhood Selection (PNS), to estimate a gene network [5]. The PNS algorithm drops genes with either 1) large p-values and/or 2) low pairwise correlation with other genes before applying the regression-based neighborhood selection method from Meinshausen and Bühlmann [8] to construct the final network. More information on the network construction procedure can be found in Supplementary Section S1.3.2, along with the various gene set sizes after filtering and within PNS (Supplementary Table S1).

Our new methodology is an extension of the DAWN algorithm, which seeks to identify sets of coexpressed genes with similar functions within a sparse network [4]. In its original formulation, the algorithm links an AP score (*Z*) with each gene in a connected network. Each gene has a hidden binary state (*I*) indicating its unobserved state (i.e., AP or not). The DAWN algorithm infers the hidden binary states of all genes in the network by leveraging the clustering of high AP scores among connected genes. As discussed in the introduction, this method assumes that AP genes tend to cluster, a theory that has been shown in numerous studies and analyses [4, 5, 6, 9, 10].

We generalize DAWN to model two sets of scores (DE and AP) jointly, within a four-state HMRF model. Our objective is to infer which of the following four hidden states each gene is most likely to be: (1) both DE and AP (active), (2) only DE (p-reactive), (3) only AP (other) and (4) neither DE nor AP (other) (Figure 2). As with the original formulation, the four-state HMRF model utilizes the scores for each measure of the gene of interest, as well as those of the neighboring genes defined by the gene network, to infer the hidden state of all genes. Similar to the original DAWN assumption, namely that AP genes tend to cluster, we assume that both AP and DE genes cluster [11]. We classify genes into their *most likely hidden states* (MLHS) according to which state has the highest posterior probability, given their observed DE and AP scores, and the gene network. All DE genes qualify as p-reactive (state 2); to change to state 1, they must cluster with many active genes. Although genes with an inferred state of 3 are also implicitly active (AP only), we label state 3 genes as “other” because they are not helpful in classifying DE genes.

**Figure 2:**
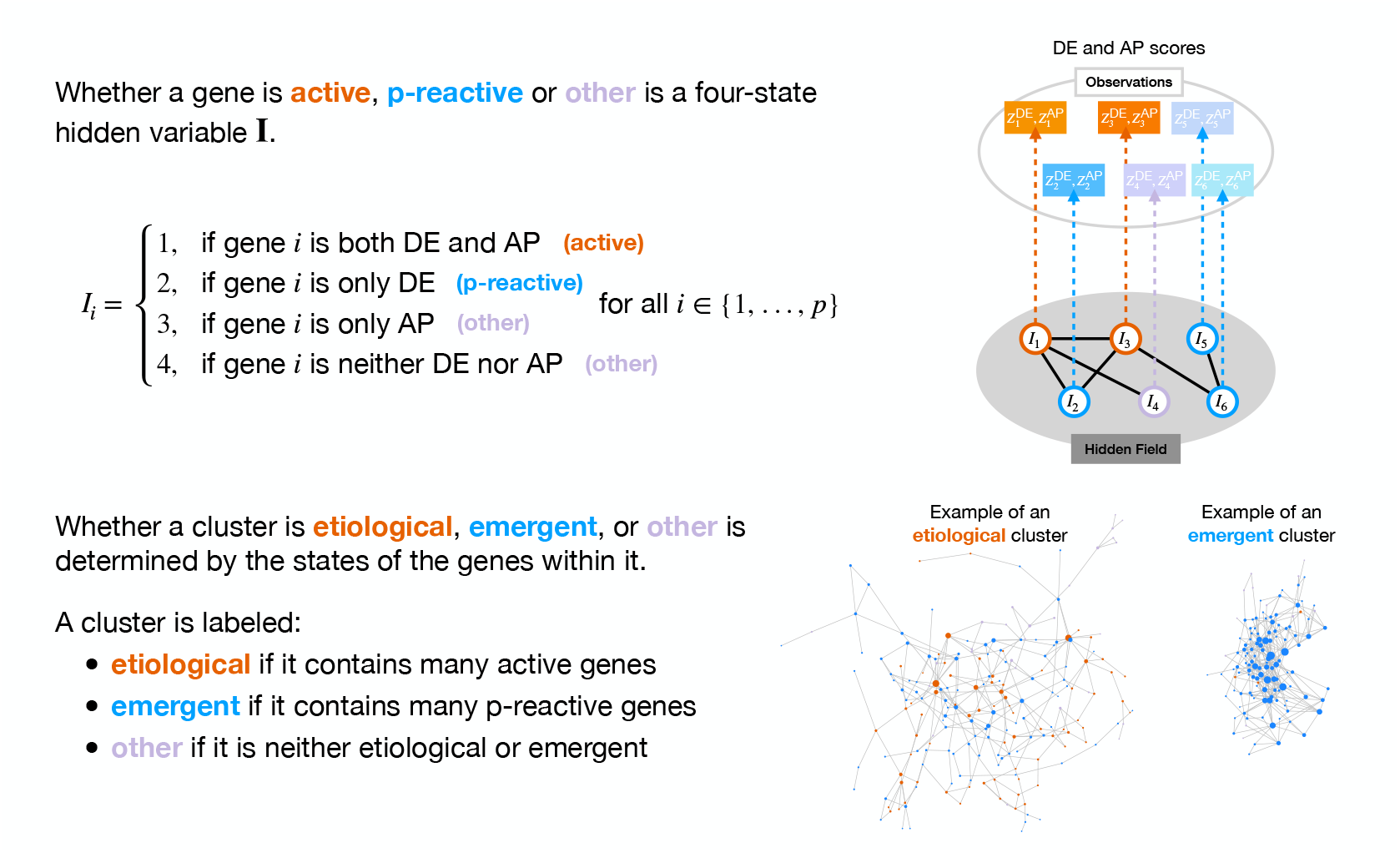
Specifics of the joint-HMRF model and cluster labeling procedure for DAWN-SCAPE. Rather than the traditional 2-state configuration for DAWN, we utilize a 4-state configuration to model the input from two sources: genetic association (AP) and differential expression (DE) scores. The unobserved, or hidden, states are superimposed on the gene network where each gene is a node. A hidden random Markov field (HMRF) model is used to infer the state of each gene. Genes are then clustered based on the network structure and labeled as etiological, emergent, or other based on the relative proportion of each state within the cluster as well as across clusters.

For clustering, we adapt the Leiden method [12], which ensures that all communities are well-connected. Despite its useful properties, the Leiden algorithm is modularity-based and does not allow the user to predefine the number of desired communities (*k*). Furthermore, the algorithm sometimes yields an overly fine-grained partition consisting of many small, highly similar communities. To address these limitations, we developed a more refined approach that allows for the re-merging of clusters and the designation of the target number of communities (*k*). We term this novel methodology Two-Step Leiden. It comprises the following sequence: (1) use Leiden [12] to construct high modularity clusters; (2) construct a similarity matrix among those clusters using information from each cluster’s low-dimensional embeddings; and, (3) using hierarchical clustering based on this similarity matrix, merge the initial clusters until we get *k* clusters (Algorithm 3).

In the network, each gene cluster is characterized by two measures of state enrichment: the proportion of each state within the cluster (internal composition or “by cluster”), and the proportion of the overall gene state represented by the cluster (global representation of that state or “by state”). Our goal is to determine which p-reactive genes are more likely to be active based on these two measures of state enrichment (Figure 2). Clusters that are enriched with an unusual number of genes in state 1 (active) are labeled as “etiological”. And clusters that are primarily state 2 (p-reactive) are labeled as “emergent”. This labeling splits gene clusters into those more likely play a fundamental role in the development of the phenotype and those differentially expressed as a consequence of the phenotype.

## Results

### ASD

ASD is a neurodevelopmental condition that arises largely from genetic variation. This variation can be transmitted over generations or appear *de novo* in the individual with ASD [13, 14, 15, 16]. Although heritable variation contributes the most to prevalence [14, 15, 16], the distribution of *de novo* exonic variation has been the most useful for associating specific genes [13]. Using the AP method to analyze *de novo* and transmitted rare variation in parent-offspring trios [17], over one hundred genes have been associated with ASD (AP genes), although more are yet to be discovered [18, 17, 13]. For AP genes, we used the results from Fu et al. [13], who reported 185 genes significantly associated with ASD (*FDR <* 0.05).

In one of the largest studies to date, Gandal et al. [1] sequenced RNA from postmortem cortical tissue of 49 ASD and and 54 neurotypical (NT) individuals. RNA was quantified from 725 samples from 11 distinct cortical regions. Using PNS and gene expression profiles from NT subjects (frontal, temporal, and occipital regions), we derived a sparse gene network. Furthermore, for the DE scores, we used the differential expression results reported in Gandal et al. [1], who identified 4 223 significant genes (*FDR <* 0.05). The distribution of AP scores appeared to be well-distributed as a mixture of 𝒩 (0, 1) and 𝒩 (*δ, τ*). Because the DE scores appeared to have a shifted null distribution, we recalibrated them using Efron’s method [19] (Supplementary Figures S1i, S2i). Taking the intersection of genes in the gene expression profiles, the AP scores, and the DE scores, we are left with 15 633 genes. For computational efficiency and to ensure relevance, we performed additional gene filtration steps based on the proportion of non-zero cells and mean expression (Supplementary Section S1.3.2).

To build a network via PNS, we chose the lasso penalty parameter *λ* = 0.22 based on a series of metrics, including visually rich structure and average degree (Supplementary Section S1.3.2, Supplementary Figure S3). After obtaining the estimated network, we pruned it further by removing disconnected genes, which will not contribute information to the network, leaving 1 218 genes. With these genes, we ran the four-state joint-HMRF to infer hidden states (active, p-reactive, or other) for all genes (Figures 2, 3a; Supplementary Table S4). Clustering via Two-Step Leiden (Supplementary Section S1.3.4) identified 14 clusters that ranged in size from 11 to 224 genes (Figure 3a-b, Supplementary Table S5). Three clusters, each with less than 20 genes, were not considered further. The 11 remaining clusters were visually apparent in the estimated network (Figure 3c).

**Figure 3:**
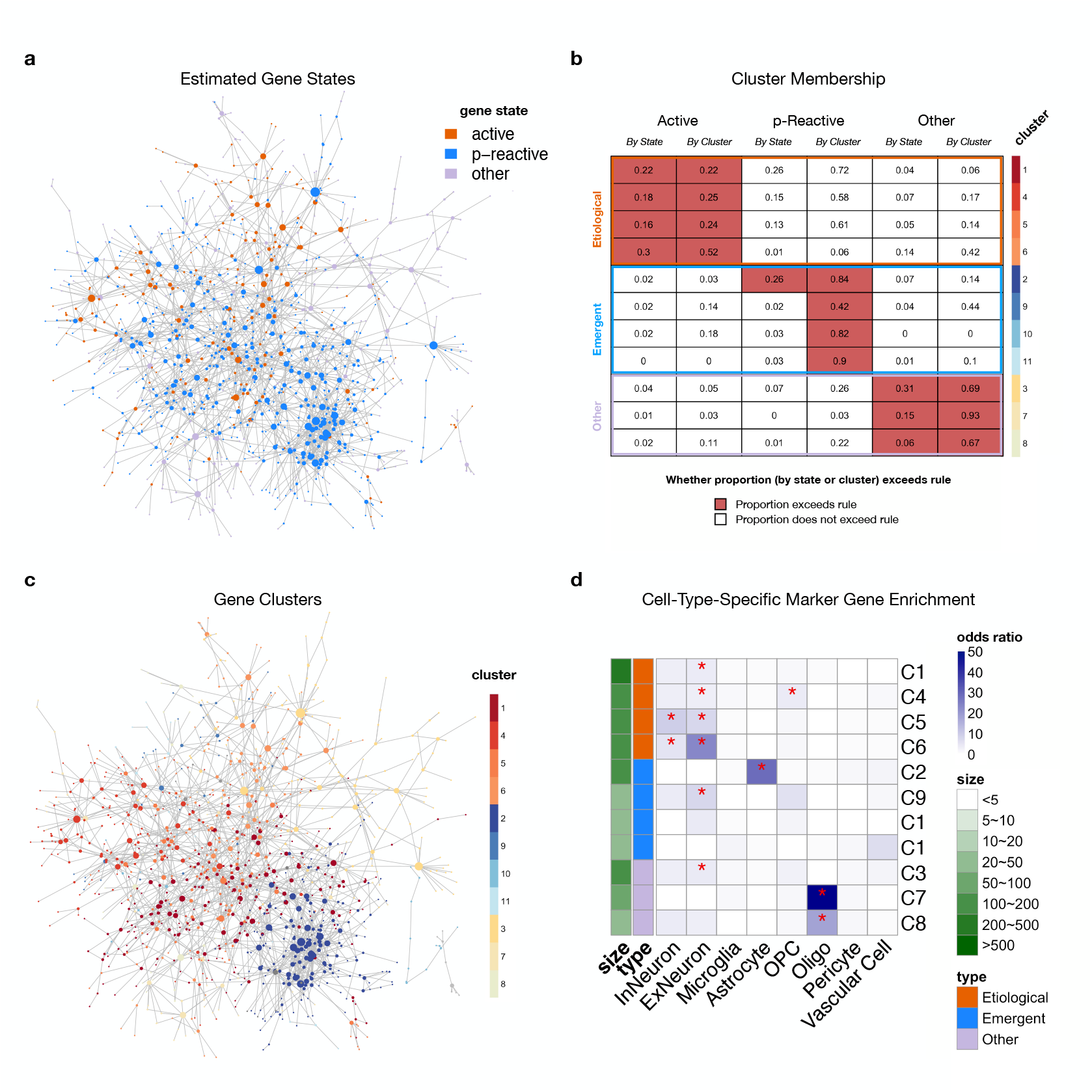
Identification of etiological, emergent, and other gene clusters in autism spectrum disorder using DAWN-SCAPE and subsequent cell-type enrichment analysis for biological context. Clusters with 20 genes or more are included in downstream analyses. (a) Gene network with nodes colored by inferred hidden state label: orange for “active”; blue for “p-reactive”; and light purple for “other”.(b) The proportion of active, p-reactive, and other genes in each of the gene clusters by state and by cluster. Cells are colored if they meet the threshold defined in Supplementary Table S2. Clusters are outlined according to their classification: etiological (orange), emergent (blue), and other (light purple). (c) Gene network with nodes colored by cluster. Genes in the three small clusters (*<* 20 genes) are colored gray and not included in the legend. (d) Heatmap displaying odds ratio for enrichment of cell-type-specific marker genes for each of seven cell types. Stars indicate significant enrichment after correcting for multiple testing (p-value cutoff = 0.05/(# tests)).

To identify etiological clusters, those enriched for active genes, versus those primarily composed of p-reactive genes (emergent), we evaluated the proportion of active and p-reactive genes in each cluster. There were clear distributional differences across clusters (Figure 3c). For example, clusters 6 and 2 were clear examples of etiological and emergent communities respectively (Figure 2): cluster 6 has many active genes, whereas cluster 2 is dominated by genes that are labeled p-reactive. Based on a set of rules involving clustering of states, clusters were classified into etiological (C1, C4, C5, C6), emergent (C2, C9, C10, C11), and other (C3, C7, C8) (Supplementary Section S1.3.5). To assign biological context to the DAWN-SCAPE clusters, we leveraged cell-type-specific marker genes derived from single-nucleus RNA-sequencing (snRNA-seq) dataset [20]. This dataset comprises 413 682 nuclei across 108 cortical tissue samples from 60 prenatal and postnatal human individuals with no known neuropathological abnormalities. To match the age range of our expression profiles for gene network construction, we subsetted this snRNA-seq dataset to include only cells derived from individuals aged 2 years and older, and combined subtypes for broader cell types (Supplementary Table S3). We first identified marker genes for the seven major cell types (interneurons, excitatory neurons, microglia, astrocytes, OPC, oligodendrocytes, pericytes, and vascular cells) using Seurat’s `FindMarkers` [21, 22] (Supplementary Section S2.1). We then tested each of the 11 clusters for cell type-specific marker gene enrichment, with the marker genes presented in Supplementary Table S6. All four etiological clusters were enriched for neuronal cell type markers, while two of the emergent clusters were enriched with CTs markers – one with excitatory neuron markers, C9, and one with astrocyte markers, C2 (Figure 3d, Supplementary Table S7). This pattern suggests that the upstream etiological processes in ASD are primarily confined to core neuronal mechanisms, whereas the emergent, downstream effects reflect a more widespread, reactive cellular environment, supporting a progression from neuronal origin to broader cellular alterations. We also compared our identified clusters with the 35 WGCNA modules from Gandal et al. [1], which are annotated with cell-type-specific enrichment information using Expression Weighted Cell Type Enrichment (EWCE) [23]. This comparison showed general congruence between the two analyses. Specifically, our etiological clusters largely overlapped with neuronal WGCNA modules. Furthermore, correspondence was observed between our excitatory neuron emergent cluster and a WGCNA module annotated with the same cell type, as well as between our astrocyte emergent cluster and a WGCNA module annotated with the same cell type (Supplementary Section S2.2, Supplementary Figure S5).

To assess biological functions of these genes, we conducted enrichment analysis using `enrichGO` function in `clusterProfiler` [24, 25], with Benjamini-Hochberg [26] for correction of multiple testing and SynGO [27] for assessing synapse-specific enrichment. We conducted two levels of analysis, cluster-specific and another in which clusters of the same type (etiological or emergent) were pooled (Supplementary Tables S8-S10). The pooled etiological gene set produced proteins affecting various synaptic and neural functions, whereas the emergent gene set largely encoded proteins that respond to stress (Figure 4). Cluster-specific analyses showed similar patterns (Supplementary Figure S7). Furthermore, a strong directional dysregulation was observed between the two gene sets: genes in the pooled etiological set were highly downregulated, while those in the emergent set were highly upregulated (Supplementary Figures S9, left panel). This opposing directional pattern by cluster type (etiological vs. emergent) is further reflected in the individual cluster log fold change (logFC) distributions (Supplementary Figure S10a-b). These findings strongly align with the long-standing hypotheses in ASD research, suggesting that initial genetic risk primarily impacts core neuronal signaling and development, while subsequent changes involve broad immune and cellular environment responses. These results were not due to our gene filtration: for the 8 000 genes we selected for analysis, the mean of logFC is −0.034 (Supplementary Figure S10c). These results also concur with previous findings: Gandal et al. [1] found that within the “Attenuation of Transcriptomic Regional Identity” genes, neuronal markers and complementary gene sets tended to be strongly down-regulated, while the upregulated genes tended to be enriched for non-neuronal cell-type markers.

**Figure 4:**
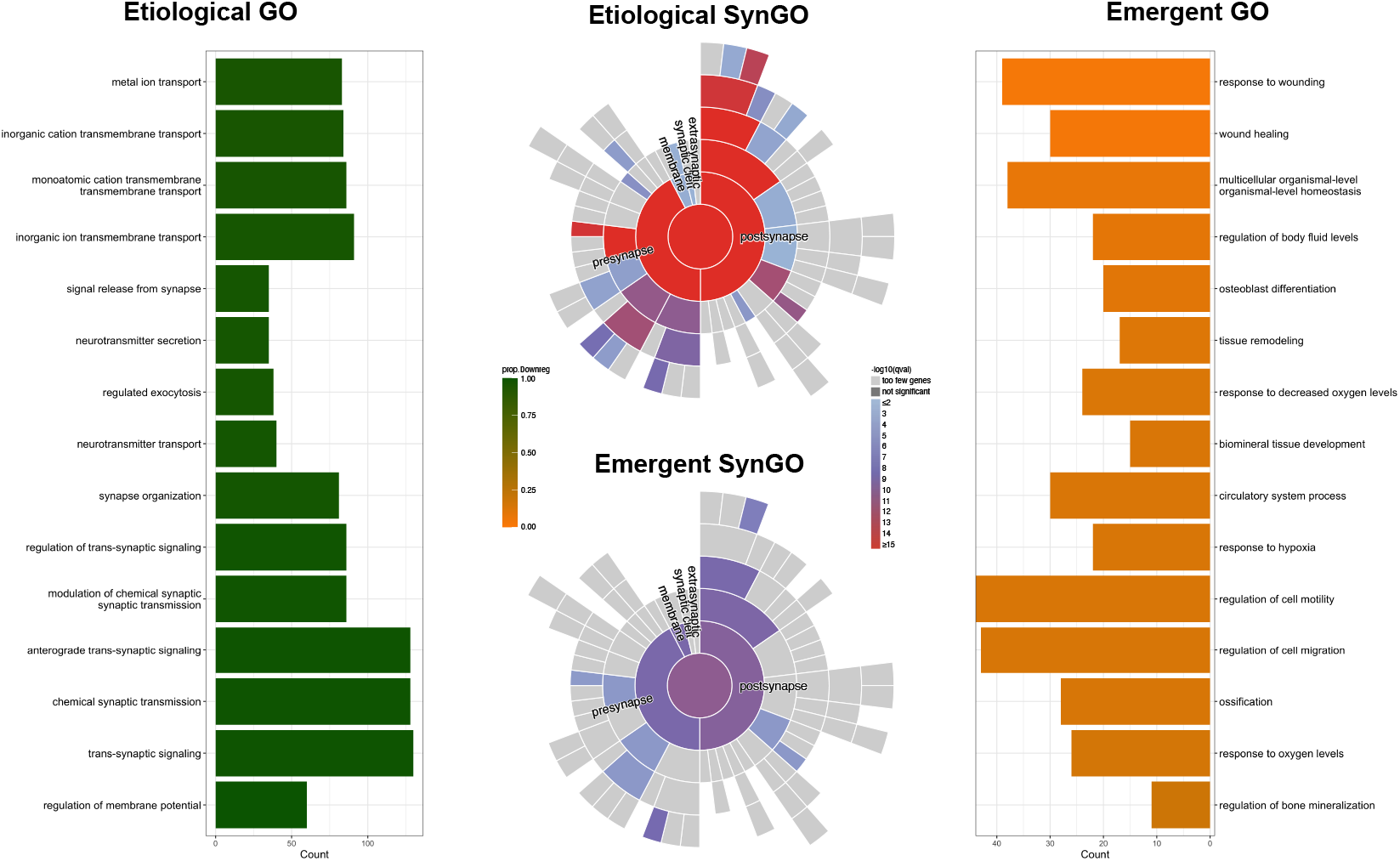
Gene ontology (GO) and synaptic GO (SynGO) enrichment results for the grouped etiological and emergent clusters for autism spectrum disorder (ASD). The top 15 GO enrichment terms by adjusted *p*-value for each set are shown. Bars are colored by the proportion of genes that are down-regulated in the GO term. The SynGO sunburst plots are colored by the −log_10_(*q*-value) for the SynGO term, where light gray indicates too few genes and dark gray indicates a non-significant *q*-value. Full GO enrichment results for the grouped clusters can be found in Supplementary Tables S8-S9. The ASD cluster-specific GO and SynGO enrichment results are presented in Supplementary Figure S7 and Supplementary Table S10.

### Alzheimer’s Disease

As the most common neurodegenerative disorder, Alzheimer’s disease (AD) affects approximately 10% of Americans over the age of 70 and is observed in older adults worldwide. Similar to ASD, AD exhibits high heritability [28]. Rare, highly penetrant genetic variants contribute to early-onset forms of the disease through dominant inheritance patterns [29]. Although genome-wide association studies (GWAS) can identify common single nucleotide polymorphisms (SNPs) associated with liability, the results alone do not implicate genes. Instead, transcription-wide association studies (TWAS), which build on GWAS data and yield gene-based test statistics, are often used. Here we use a TWAS study from Liu et al. [30], who analyzed gene expression data from hippocampal tissue samples and recent AD GWAS results. From their “replication sample”, they identified 45 genes significantly associated with AD (*FDR <* 0.05) and provided *p*-values for 14 552 protein-coding genes. These results will serve as the core of the AP scores for our DAWN-SCAPE analysis. In addition to these data, we also forced into the analysis a set of genes believed to influence the liability to Alzheimer’s disease through common ([31], their Fig. 2) or rare variation ([32], their Table 1). Joining these two resources yielded 53 genes (Supplementary Table S11), which we call the fixed gene set and include genes such as *APOE* (common variation) and *PSEN1* (rare variation). After adding in the fixed gene set to the core AP scores from the TWAS, we have 97 genes significantly associated with AD.

The differences in gene expression between AD and control brain tissue are widespread and have been extensively studied. For example, a recent meta-analysis of 584 subjects and 778 tissue samples [33] identified a substantial transcriptional dysregulation, reporting 2 355 significantly up-regulated and 2 130 significantly down-regulated transcripts (*FDR <* 0.05). For our DAWNSCAPE analysis, we utilize the *p*-values for 13 103 protein-coding genes from this meta-analysis’ random effects model as our DE scores (SYN11914808). For the gene network, we used the gene expression profiles from Gandal et al. [1]. Taking the intersection of genes in the gene expression profiles, the AP scores, and the DE scores, we are left with 11 772 genes.

The AP and DE scores were prepared for network analysis using the same processing pipeline as described for ASD. Briefly, this involved checking the AP and DE scores and recalibrating the DE scores using Efron’s method due to a shifted null distribution (Supplementary Figures S1ii, S2ii), followed by gene filtration based on non-zero cell proportion and mean gene expression. The network was then constructed using the PNS approach, employing Meinshausen and Bühlmann [8]’s LASSO-regression method. We chose the lasso penalty parameter *λ* = 0.16 for PNS based on a series of metrics, including visually rich structure and average degree (Supplementary Section S1.3.2, Supplementary Figure S4). After obtaining the estimated network, we pruned it further by removing singleton genes, leaving 2 113 genes. With these genes, we ran four-state joint-HMRF to classify genes as active, p-reactive or other (Figures 2, 5a; Supplementary Table S11).

**Figure 5:**
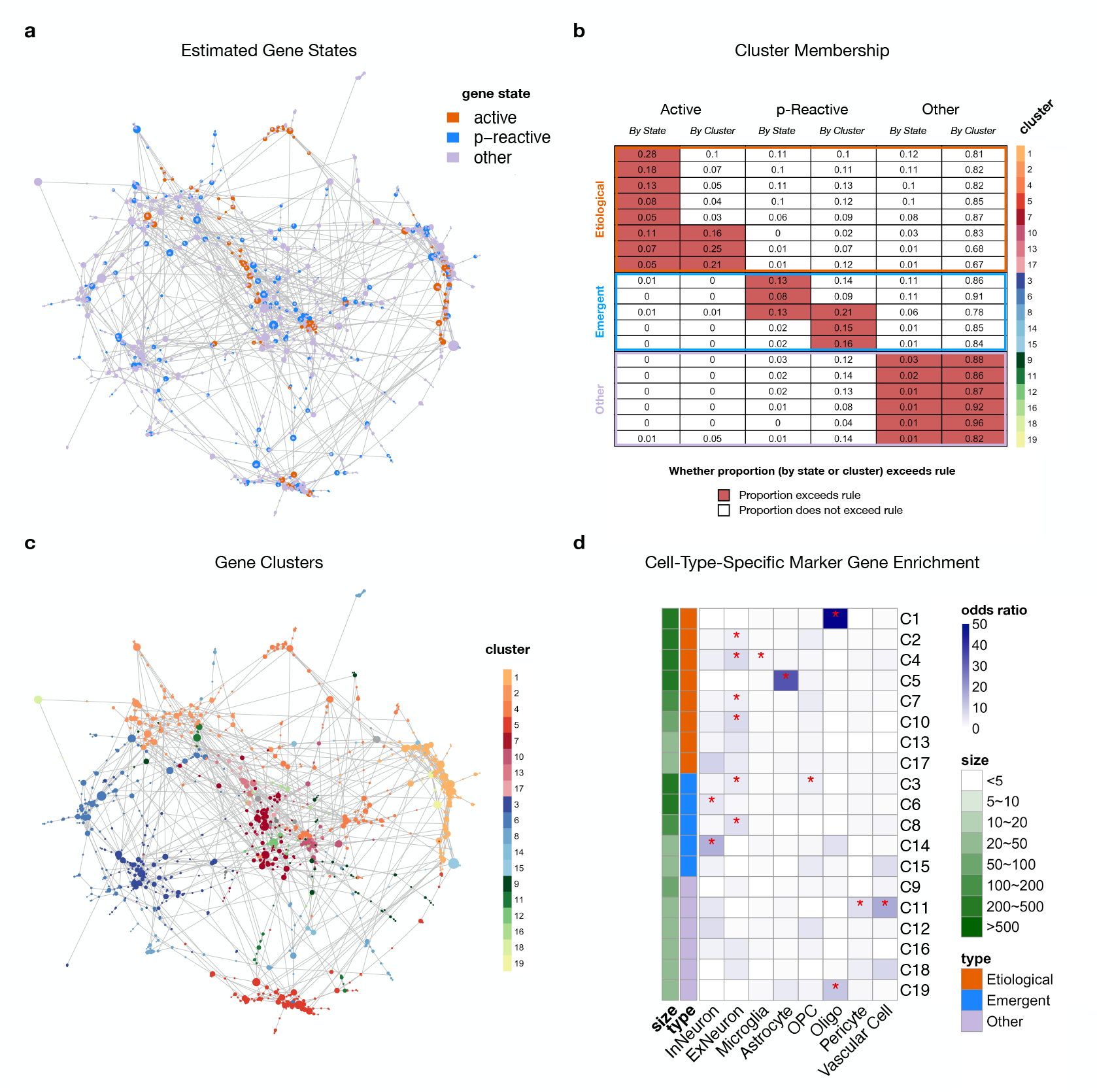
Identification of etiological, emergent, and other gene clusters in Alzheimer’s disease using DAWN-SCAPE and subsequent cell-type enrichment analysis for biological context. Only clusters with 20 genes or more are included in downstream analyses. (a) Gene network with nodes colored by inferred hidden state label: orange for “active”; blue for “p-reactive”; and light purple for “other”. (b) The proportion of active, p-reactive, and other genes in each of the gene clusters by state and by cluster. Cells are colored if they meet the threshold defined in Supplementary Table S2. Clusters are outlined according to their classification: etiological (orange), emergent (blue), and other (light purple). (c) Gene network with nodes colored by cluster. Genes in the three small clusters (*<* 20 genes) are colored gray and not included in the legend. (d) Heatmap displaying odds ratio for enrichment of cell-type-specific marker genes for each of seven cell types. Stars indicate significant enrichment after correcting for multiple testing (p-value cutoff = 0.05/(# tests)).

Our clustering procedure (Section S1.3.4) identified 23 clusters that ranged from 10 to 272 genes (Figure 5b, Supplementary Table S12). Four clusters, each with less than 20 genes, were not considered further. The 19 remaining clusters were visually apparent in the estimated network (Figure 5c). Consistent with the relatively small number of active genes entering the analysis, none of the clusters have many active genes. Nonetheless, it was still possible to classify the 19 clusters into “etiological”(C1, C2, C4, C5, C7, C10, C13, C17), which have higher proportions of active genes; “emergent” (C3, C6, C8, C14, C15), with a greater fraction of p-reactive genes; and “other” (C9, C11, C12, C16, C18, C19) (Supplementary Table S5). As in ASD, we conducted a cell type-specific marker gene enrichment analysis to assign a biological context to our clusters, with the full list of marker genes provided in Supplementary Table S13. Interestingly, astrocyte markers were strongly enriched in one etiological cluster, C5, whereas oligodendrocyte markers were enriched in another, C1 (Figure 5d, Supplementary Table S14). This aligns with increasing evidence that astrocytes and oligodendrocytes are fundamental to early etiological stages of AD [34], supporting roles in synaptic clearance failure and neuroinflammation [35, 36] as well as white matter degeneration [37, 38]. Emergent clusters were mostly enriched in neuronal cell type markers, with the one exception being OPC markers in C3. Comparing our clusters with the 35 WGCNA modules from Gandal et al. [1], we see a high similarity between our etiological and emergent clusters and the neuronal WGCNA clusters (Supplementary Section S2.2, Supplementary Figure S6). Additionally, we find a large overlap between our astrocyte etiological cluster and a WGCNA module annotated with the same cell type, as well as between our oligodendrocyte etiological cluster and a WGCNA module annotated with the same cell type.

As with ASD, we conducted both a cluster-specific analysis and a pooled-cluster analysis. GO enrichment and quantitative differential expression (logFC) analyses reveal modest overall upregulation in the pooled etiological gene set, whereas the pooled emergent gene set was distinctly downregulated (Figure 6, Supplementary Figure S9, right panel). While many of the genes in both the pooled etiological and emergent sets were involved in neuronal and synaptic functions, their expression tended to be strongly downregulated, consistent with neurodegeneration (Supplementary Tables S15-S16). Additionally, within the etiological set, genes associated with gliogenesis were upregulated, aligning with the widely-held theory that activated microglia and astrocytes drive neuroinflammation and subsequent neurodegeneration in AD [35, 39], with more recent research highlighting the active contribution of oligodendrocytes to this inflammatory process [34, 38]. This pattern of upregulation persisted for other significant functions related to glial cells, as evidenced by the GO terms emerging from enrichment analysis. Other genes in the etiological set functioned in vascular systems, although there was no consistent direction of their expression. Nonetheless, this enrichment aligned with the vascular theory of AD, which posits a relationship between cardiovascular pathology and AD-related changes [40]. While the exact nature of this relationship is still debated, our results suggest an upstream association with AD and align with a core tenet of the hypothesis, namely that cardiovascular disease is either causal or co-occurring with AD pathology.

**Figure 6:**
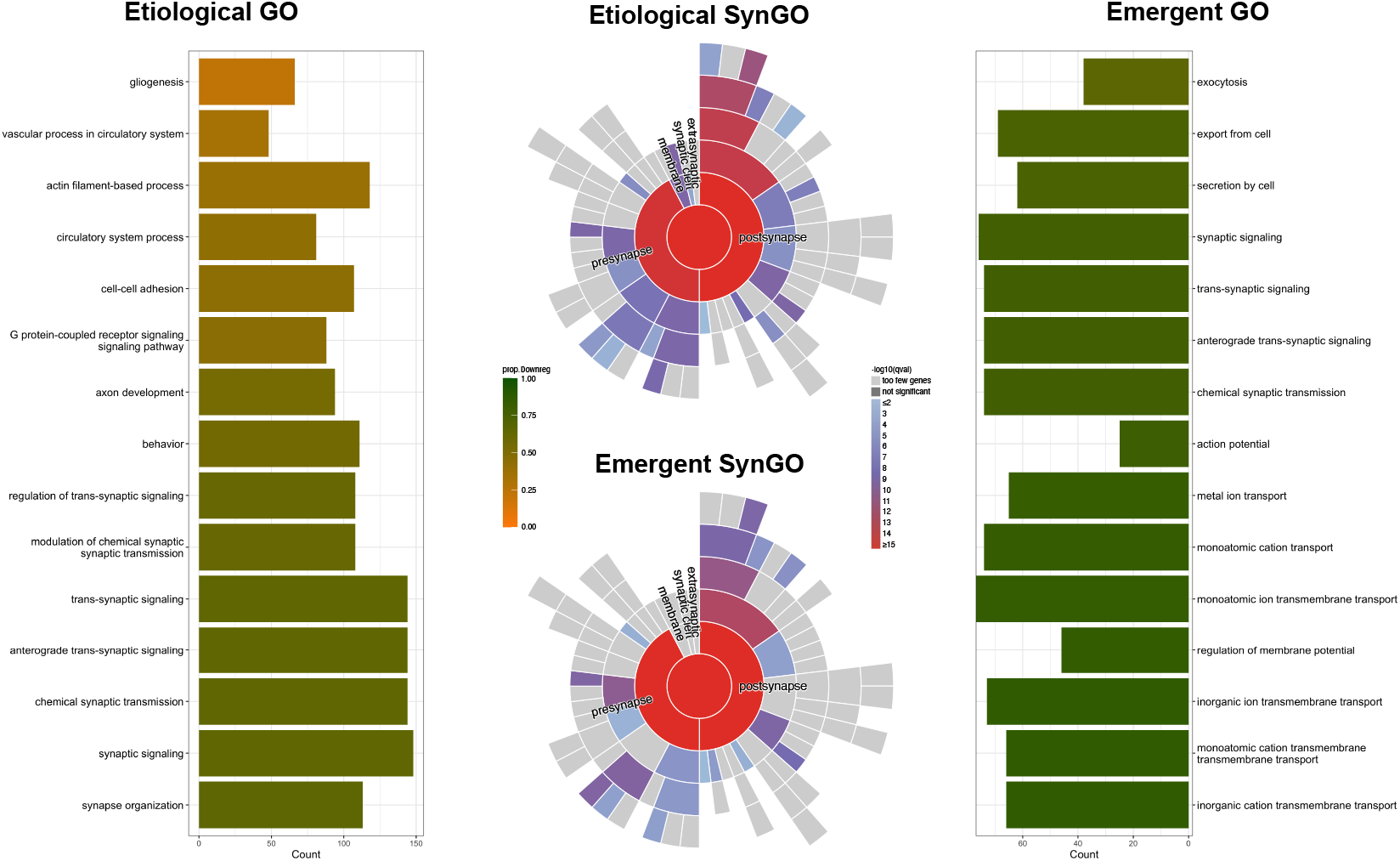
Gene ontology (GO) and synaptic GO (SynGO) enrichment results for the grouped etiological and emergent clusters for Alzheimer’s disease (AD). The top 15 GO enrichment terms by adjusted *p*-value for each set are shown. Bars are colored by the proportion of genes that are down-regulated in the GO term. The SynGO sunburst plots are colored by the −log_10_(*q*-value) for the SynGO term, where light gray indicates too few genes and dark gray indicates a non-significant *q*-value. Full GO and SynGO enrichment results for the grouped clusters can be found in Supplementary Tables S15-S16. The AD cluster-specific GO and SynGO enrichment results are presented in Supplementary Figure S8 and Supplementary Table S17.

Cluster-specific GO analysis found few etiological and emergent clusters enriched for neuronal and synaptic functions (etiological: C2, C7, C10; emergent: C6). Additionally, etiological cluster C1 was enriched for glial and axon-related functions, while etiological cluster C5 had numerous genes with glial and vascular/circulatory system-related functions (Supplementary Figure S8, Supplementary Table S17). Genes in clusters C1 and C5 were notably upregulated, particularly compared to other clusters (both etiological and emergent, Supplementary Figure S11a-b). Thus, enrichment and differential expression results were broadly consistent with current understanding of Alzheimer’s disease, such as the role of glial activation and cardiovascular systems in AD pathology.

## Discussion

By contrasting gene expression profiles of relevant tissue or cells taken from individuals of different phenotypes, such as affected and unaffected by disease, we can identify sets of genes whose expressions are dysregulated. What we cannot know from these results, however, is which genes differ because they influence the phenotype and which differ as a result of the phenotype. We develop a formal model to distinguish between genes that cause the phenotype, dubbed active genes, and those show differential expression due to the phenotypic differences (reactive genes). Our approach uses three sources of information: gene-specific association with phenotype (AP); differential gene expression (DE, or something equivalent); and gene-based networks. Building on earlier work [4, 5, 6], we develop a hidden random Markov field Model to accomplish the goal of identifying active and reactive genes. Furthermore, by assuming that there exist gene communities enriched only with DE-significant genes, and others that are enriched with both DE and AP genes, we build an algorithm to identify these emergent versus etiological communities (Figure 1). When we evaluate this algorithm using data relevant to autism spectrum disorder (ASD) and Alzheimer’s disease (AD), our results align with existing hypotheses about the origins and effects of these phenotypes.

Our method assumes that genes in the same state are proximate in the network typology. Specifically, we assume such genes are conditionally dependent, because the underlying network topology is inferred using ℓ_1_-regularized regression (LASSO). Nonetheless, empirical evidence often supports the clustering and co-expression of both AP and DE genes separately in various biological networks, and our own results support the assumption as well. A secondary constraint is the model’s dependency on a shared statistical signal between the AP and DE scores. Limited congruence between the two metrics would severely diminish the model’s effectiveness. Because differential expression analyses typically identify numerous significant genes, the primary uncertainty lies in the smaller set of genes flagged by the AP scores. Consequently, the joint-HMRF model depends on the quality and strength of the genetic results; without a shared signal, the model could fail.

A compelling direction for future work involves incorporation of covariates into the hidden random Markov field (HMRF) framework. While covariates could be integrated during network construction or the clustering process, the most informative approach would be to integrate them directly into the latent state definition. For instance, incorporating the directionality of expression change (up- or down-regulation) could yield greater mechanistic insight, which is highlighted by our results for Alzheimer’s Disease. This modification would replace the binary DE with two distinct latent state questions: (1) whether the gene is DE and up-regulated, and (2) whether the gene is DE and down-regulated, thereby increasing model complexity to eight states. Such an approach, while computationally challenging, could deliver more granular, mechanistic insights for some phenotypes.

An intriguing subset of proteins identified within the AD etiological group are those involved in actin filament processes. About a decade ago, Bamburg and Bernstein [41] hypothesized that cofilin/actin biochemistry, which impacts the dynamics of cellular filaments and thus synaptic plas-ticity, is a mechanism that could unify and explain competing hypotheses about the etiology of AD. As they note, increased abundance of active cofilin (dephosphorylated cofilin) is a pathway for synaptic loss, the hallmark of AD, and it is an expected outcome for all known mechanisms of AD (accumulation of amyloid-*β*, neurofibrillary tangles, and chronic neuroinflammation). Since then, evidence has been accumulated to support their hypothesis [42, 43, 44], and it is intriguing that our results also support this mechanism as an etiological component of the pathogenesis of AD.

## S1 Supplementary Methods

### S1.1 Algorithm Details

#### S1.1.1 Partial Neighborhood Selection

Here we describe the partial neighborhood selection method for network estimation from [5]. The full algorithm can be found in Algorithm 1. To ensure precise communication regarding the method, we have introduced specific nomenclature (e.g., the key gene set, the core gene set) to refer distinctly to the various gene sets throughout the method.

##### Algorithm 1 PNS algorithm

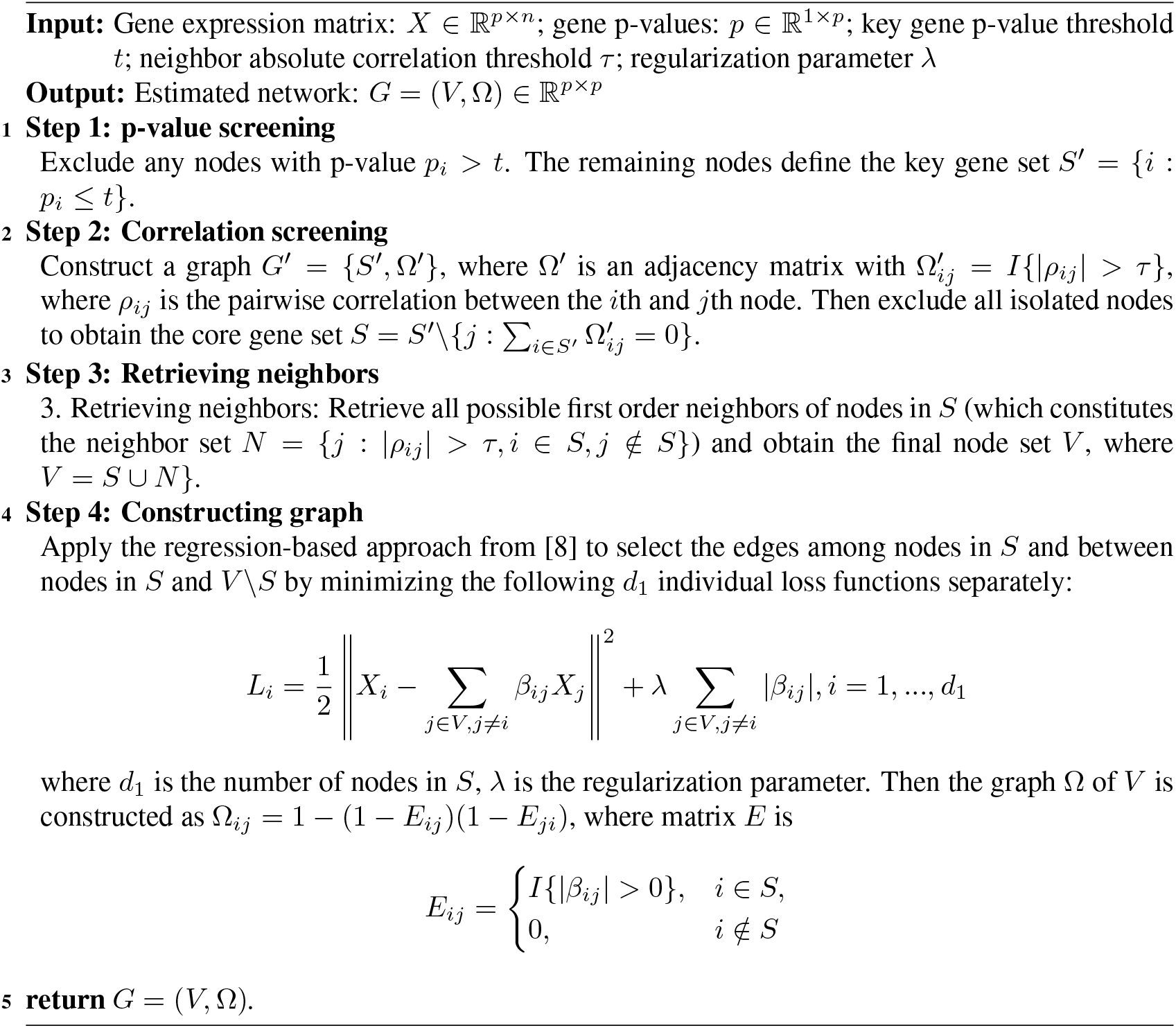

PNS takes in a gene expression matrix (bulk or single-cell); a list of p-values (e.g. corresponding to AP), one for each gene in such matrix; two thresholding hyperparameters *t* and *τ*; and regularization parameter *λ*. The PNS algorithm begins by dropping genes with p-value ≥*t*. The remaining set of genes is the key gene set, *S*^*′*^ = {*i* : *p*_*i*_ ≤*t*}(Step 1). From the key gene set, the algorithm constructs an adjacency matrix 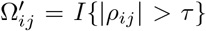where *ρ*_*ij*_ is the pairwise correlation between the *i*th and *j*th nodes, calculated using the gene expression matrix. Then, all isolated nodes, i.e. nodes with no edges, are dropped leaving the core gene set *S* (Step 2). The neighbor gene set *N* is defined as the set of non-*S* genes that have an absolute correlation greater than the threshold *τ* with at least one gene in the core gene set *S*: *N* = {*j* : |*ρ*_*ij*_|*> τ, i* ∈ *S, j* ∉ *S*}. The final node set, *V*, is then constructed by taking the union of the core gene set and the neighbor gene set: *V* ⋃ *N* (Step 3). To construct the final graph, the algorithm applies the regression-based approach from [8] to select the edges among nodes in *S* and between nodes in *S* and *V*\*S* by minimizing *d*_1_ = |*S*| individual loss functions. The final graph Ω of *V* is defined as Ω_*ij*_ = 1 − (1 −*E*_*ij*_)(1 − *E*_*ji*_) where *E*_*ij*_ = *I* {|*β*_*ij*_| *>* 0} if gene *i* is in the core set, *S*, and *E*_*ij*_ = 0 if gene *i* is just in the neighbor set, *N* (Step 4).

#### S1.1.2 Hidden Markov Random Field Model

##### Four-State HMRF Model Specification

Consider a set of *p* genes indexed by 1, …, *p*. Let 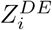 and 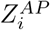 denote the observed DE and AP *z*-score respectively for gene *i*. Each gene *i* is believed to have a true, underlying hidden state category *I*_*i*_. These hidden states are assumed to follow a HMRF structure and have four possible values:

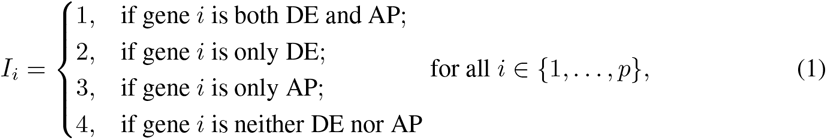

where a value of 1 or 2 indicates a gene is “active” or “p-reactive” respectively, and a value of 3 or 4 indicates “other” genes of less interest.

We then model ***Z***^*AP*^, ***Z***^*DE*^, and ***I*** using the following HMRF model. The probabilistic nature of 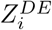 and 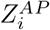 is determined by the unobservable Markov random field on {*I*_*i*_, *i* ∈ [*p*]}. That is, given the neighbors *N*_*i*_ of *i, I*_*i*_ is independent of all other *I*_*j*_ (Markov property). The model is formulated in such a way that conditioning on *I*_*i*_, 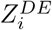, and 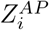 are independent of any other observable variables. We also assume that 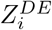 are independent of 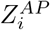.

Therefore, the joint probability of ***Z***^*DE*^, ***Z***^*AP*^, ***I*** can be written as

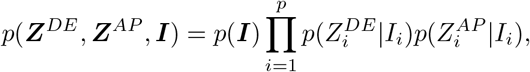

and parameters can be estimated by iteratively maximizing the three parts: 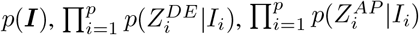.

To model 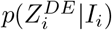, following [4], we assume each 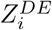 follows a Gaussian mixture model:

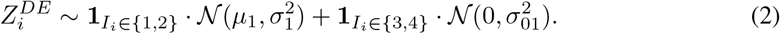

Similarly, we assume each 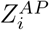 follows a different Gaussian mixture model:

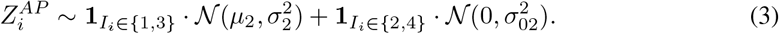

To model *p*(***I***), we design the potential function in the Markov random field such that *p*(***I***) has the following representation:

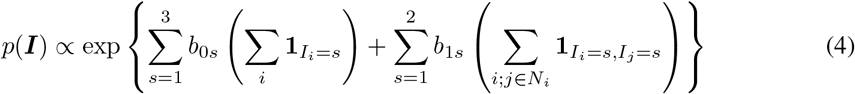

However, for quick optimization, we instead optimize the pseudo-likelihood

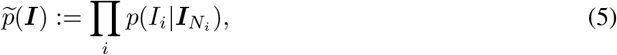

where 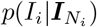 takes the form

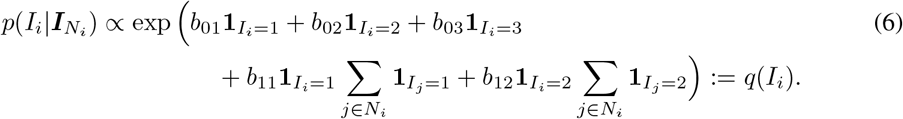

Note that the network structure is only relevant for active or p-reactive cases, as those are the only two we conjecture are clustered together meaningfully in the network.

##### Optimization

For optimization, we follow an EM style approach: alternating between 1) maximizing the pseudo-likelihood 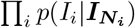 by using coordinate descent to obtain the updated interaction parameters 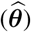, 2) estimating the maximum a posteriori (MAP) probability of the hidden states (***I***) using the iterative conditional mode (ICM) method [45], and 3) maximizing 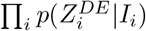 and 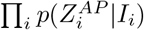) using the relationship between the Gaussian MLE and moment estimation to obtain the updated observation parameters 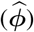.

As for initialization, instead of random initialization, we initialize the hidden states of the node using

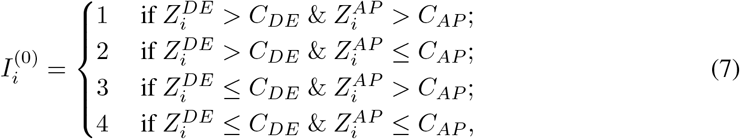

where the cutoffs *C*_*DE*_ and *C*_*AP*_ are pre-determined thresholds for the DE and AP z-scores respectively; and initialize the values of the parameters using

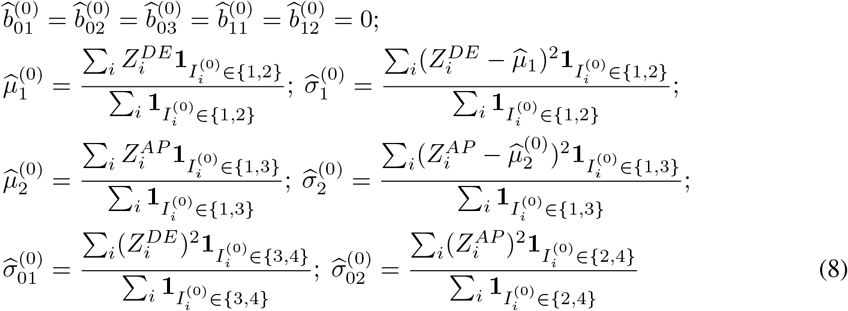

Then the whole optimization procedure is summarized in Algorithm 2. After obtaining 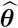 and 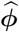, we apply a final cycle of ICM to obtain our final estimates of the hidden states, ***I***^(*t*+1)^.

###### Algorithm 2 Joint-HMRF inference

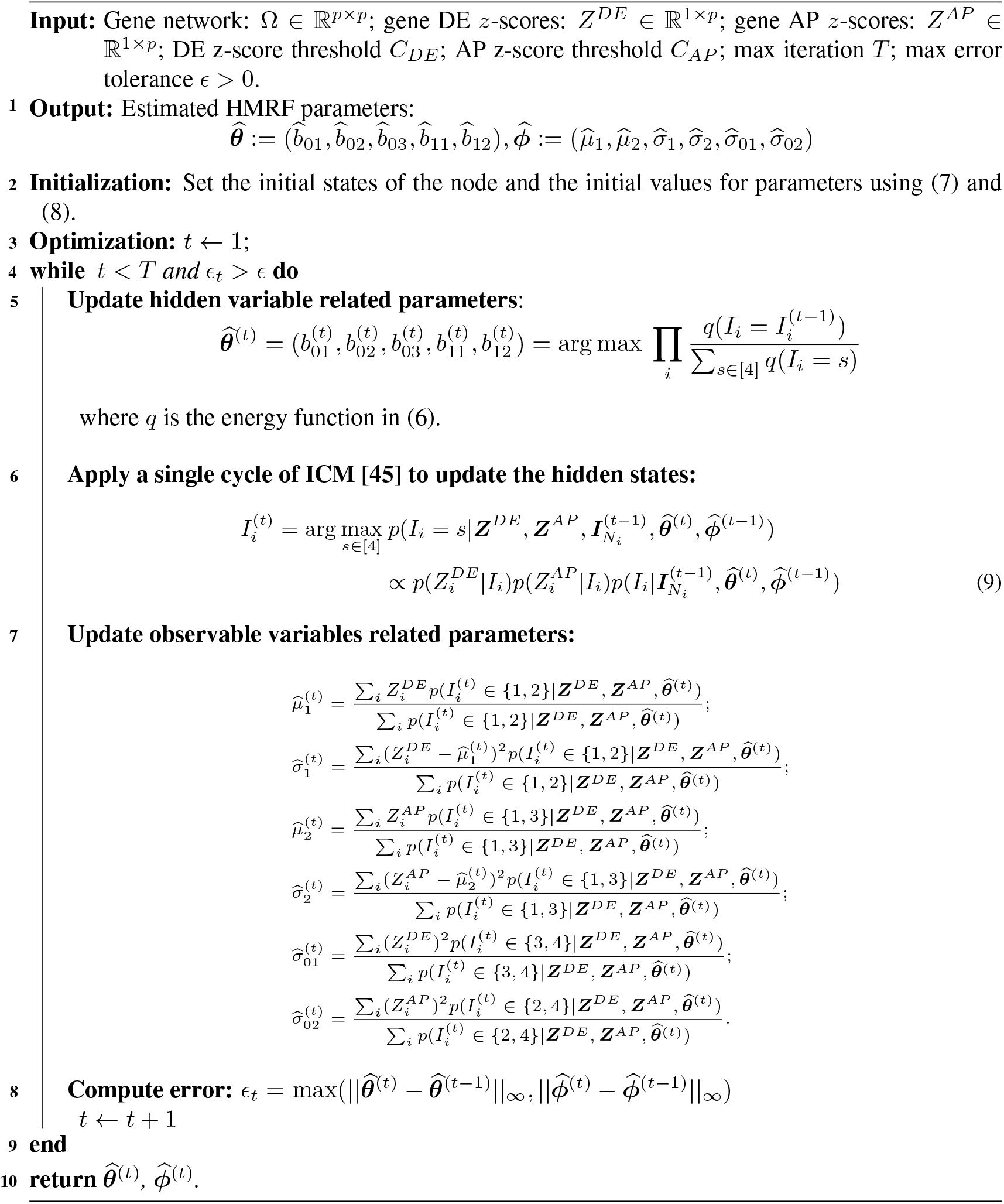

#### S1.1.3 Two-Step Leiden

At the core of our clustering procedure is an extended version of Leiden clustering [12] which we call Two-Step Leiden. We explain Two-Step Leiden in this section and an explanation of our full clustering procedure can be found in Section S1.3.4. For our data, we noticed in initial explorations that, when Leiden was run with a resolution that gave a high modularity, many of the outputted clusters tended to be too small and very similar to each other. These observations inspired the following two-step approach. We begin by using Leiden with resolution *r* to construct *l* initial clusters. From these we construct a similarity matrix using information from each cluster’s low-dimensional embedding. e.g. the first principal component. Second, we conduct a hierarchical clustering `h` based on this similarity matrix. If the number of initial clusters *l* is larger than the desired number of clusters *k*, cut `h` to merge the initial clusters and obtain *k* final clusters. The detailed steps are presented in Algorithm 3.

##### Algorithm 3 Two-Step Leiden

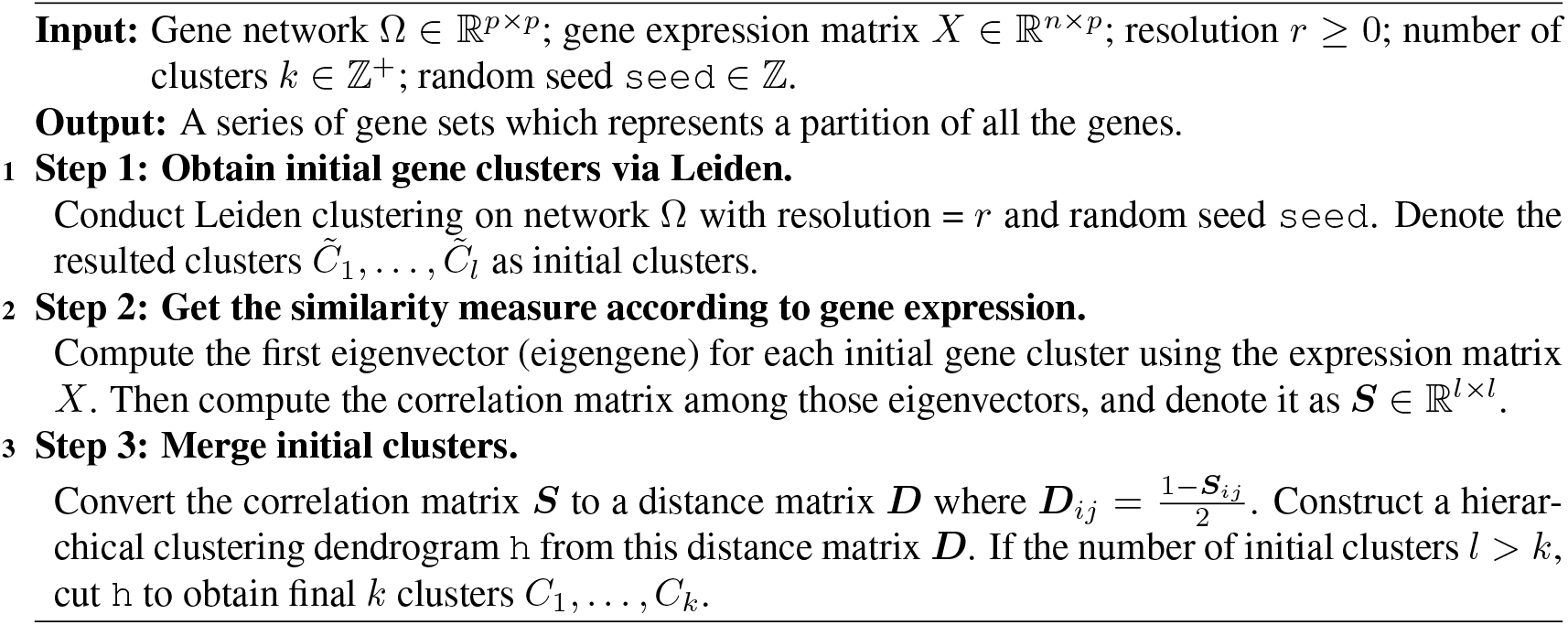

### S1.2 Description of Datasets

#### S1.2.1 Data summary

For each application, our input data is made up of three components:

##### Gene expression data for network construction

Both applications use whole cortex gene expression data (bulk RNA-sequencing data) from Gandal et al. [1] comprising samples from 54 neurotypical control individuals. For ASD we include samples from frontal, temporal, and occipital and for AD we include samples from frontal, parietal, temporal, and occipital.

##### DE significance metrics

*p*- or *q*-values obtained from differential expression analyses comparing case and control individuals. For the ASD application, *q*-values are from Gandal et al. [1] and for the AD application, *p*-values are from Fu et al. [13].

##### AP significance metrics

*p*- or *q*-values obtained from a genetic analysis such as AP or TWAS. For the ASD application, *p*-values are from Wan et al. [33]. For the AD application, the significance set is obtained by combining *p*-values from Liu et al. [30] with fixed genes believed to influence the liability to AD through common [31] or rare variation [32].

#### S1.2.2 Gene Expression Data for Network Construction

Gandal et al. [1] perform bulk RNA-sequencing (RNA-seq) on 725 brain samples spanning 11 distinct cortical areas in 112 ASD cases and neurotypical controls. The authors have conducted several processing and analysis steps shown below, which prepare us for our analysis.

1. `Gene Filtering`: Genes were retained if they had a counts-per-million (CPM) value greater than 0.1 in at least 30% of the samples. Additionally, genes with an effective length (measured by RSEM) of less than 15 bp were removed. Following these filters, the dataset consisted of 24 836 genes.
2. `Normalization`: To ensure comparability and eliminate potential biases, the remaining genes were subjected to further normalization using the limma-trend approach within the `limma` R package. This approach involved taking the log2(CPM+1) transformation of read counts, while accounting for variations in sample read depth. Additionally, a CQN-derived offset value was incorporated during the normalization process to address potential biases related to GC content and gene effective length. Collectively, these steps aimed to obtain normalized expression data suitable for downstream analysis.
3. `Outlier removal`: To identify sample outliers within each sequencing batch by cortical lobe group (frontal, parietal, temporal, and occipital), the normalized expression data underwent a two-step outlier detection process. First, samples were flagged as outliers if they met the following criteria: (1) an absolute z-score exceeding 3 for any of the top 10 expression principal components (PCs), and (2) a sample connectivity score below −2. The sample connectivity score was computed using the fundamental `NetworkConcepts` function from the `WGCNA` R package, utilizing the signed adjacency matrix (soft power of 2) of the sample biweight midcorrelation. This procedure successfully identified 34 outliers in the dataset.
4. `Technical effects removal`: Next, to address technical effects, a regressed dataset was created using the `lmerTest` package in R. This involved subtracting the effects of 20 technical covariates from each gene, resulting in a dataset that retained only the random intercept, biological covariate effects, and the residual. The regressed gene expression dataset specifically captured the effects of biological covariates, including `subject`, `diagnosis`, `region`, `sex`, `ancestry`, and `age`.

The above processing output a final gene expression matrix with a total of 24 836 genes and 725 samples (341=control, 384=ASD) with technical noises removed. We use the control samples for network construction. This gene expression matrix was also used by the authors to construct WGCNA modules which we compare against later in the Supplement. Additionally, the authors used `limma` to model cortex-wide differential expression, obtaining logFC and *q*-values. This process included using 1) `duplicateCorrelation` to account for repeated measures by subject across multiple brain regions, 2) fitting a linear model while controlling for known biological and sequencing related technical covariates including `diagnosis`, `region`, `sequencing batch`, and `sex` which produced logFC and unadjusted *p*-values, and 3) using `eBayes` and `topTable` to compute FDR-adjusted *p*-values.

### S1.3 Details of Pipeline

#### S1.3.1 Recalibration

Assessment of the distributional properties of both sets of measures of statistical significance, DE and AP, was conducted to validate their use in the subsequent model. When we have *q*-values rather than *p*-values, we first transform the *q*-values into *p*-values using the following formula for each gene *i* ∈ [*p*]:

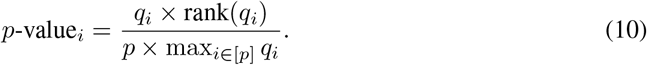

We then check whether the *p*-values require re-calibration. Specifically, we consider the *p*-values well-calibrated if the distribution of the corresponding *z*-scores using

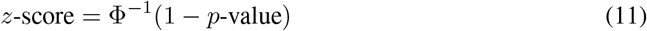

fits a mixture of *N* (0, 1) (null) and *N* (*δ, τ*) (signal), with the null signal being the predominant component.

**Figure S1:**
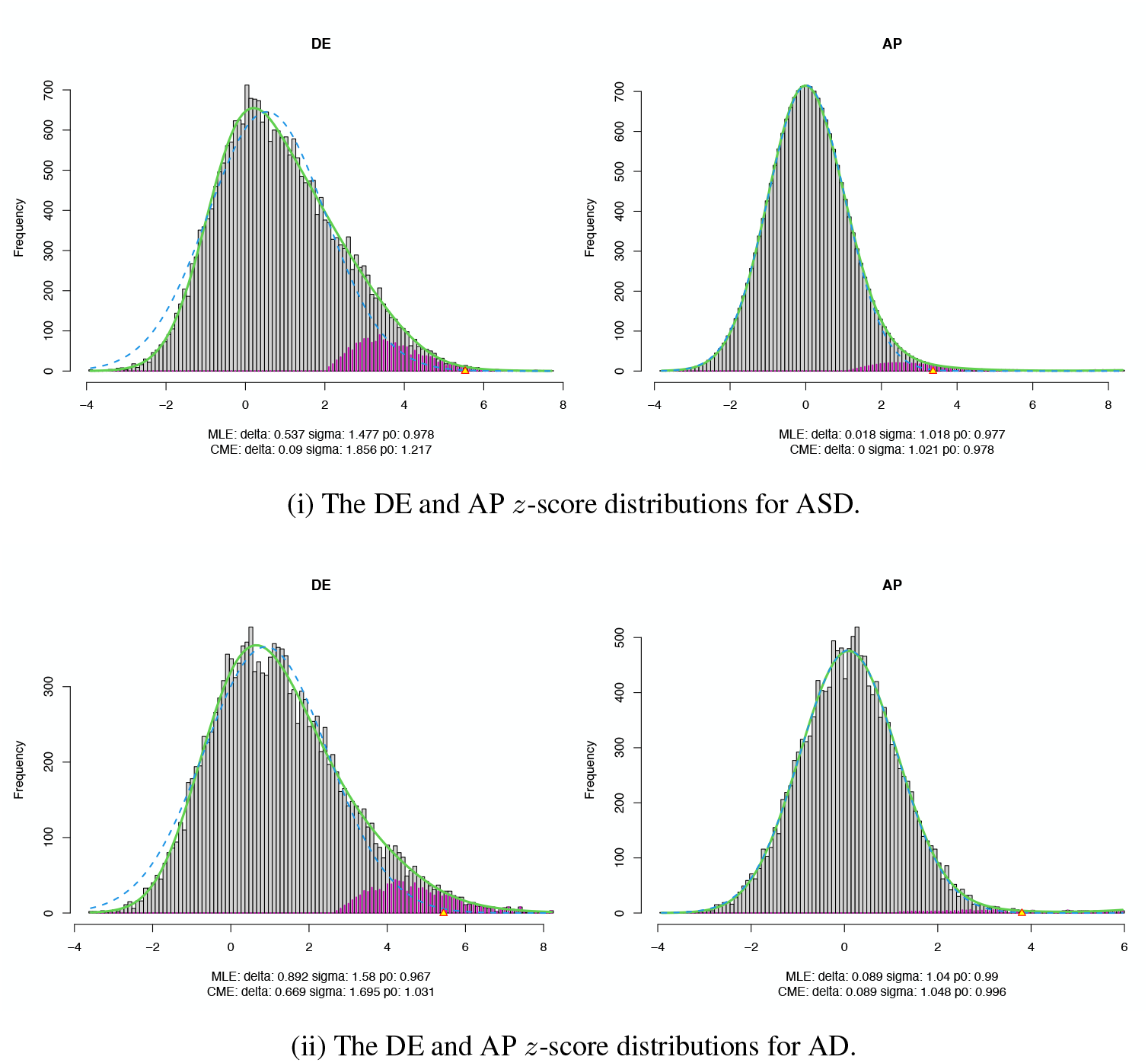
The DE and AP z-score distributions for (i) autism spectrum disorder (ASD) and (ii) Alzheimer’s disease (AD) modeled as a mixture of two underlying components, the null (grey) and non-null or signal (purple) distributions, using Efron’s `locfdr` method [19].

However, before we check the *z*-score distribution, we need to perform a couple of operations on the *p*-values and *z*-scores. As discussed in the main paper, for AD, we identified 53 genes thought to influence liability to AD through common or rare variation. We call these genes “fixed” genes. We forced the fixed genes into the analysis using the following procedure: As long as the gene existed in our bulk gene expression dataset for network construction, we were able to include it in our analysis. For these genes, we set the AP *p*-value to be 0.0 and, if the gene didn’t exist in the DE *p*-value set, we set it to be 0.5, a non-significant, neutral placeholder. Ultimately, 52 of the 53 genes were part of the bulk gene expression dataset and were included in the 8 000 genes considered for network construction in Supplementary Section S1.3.2.

**Figure S2:**
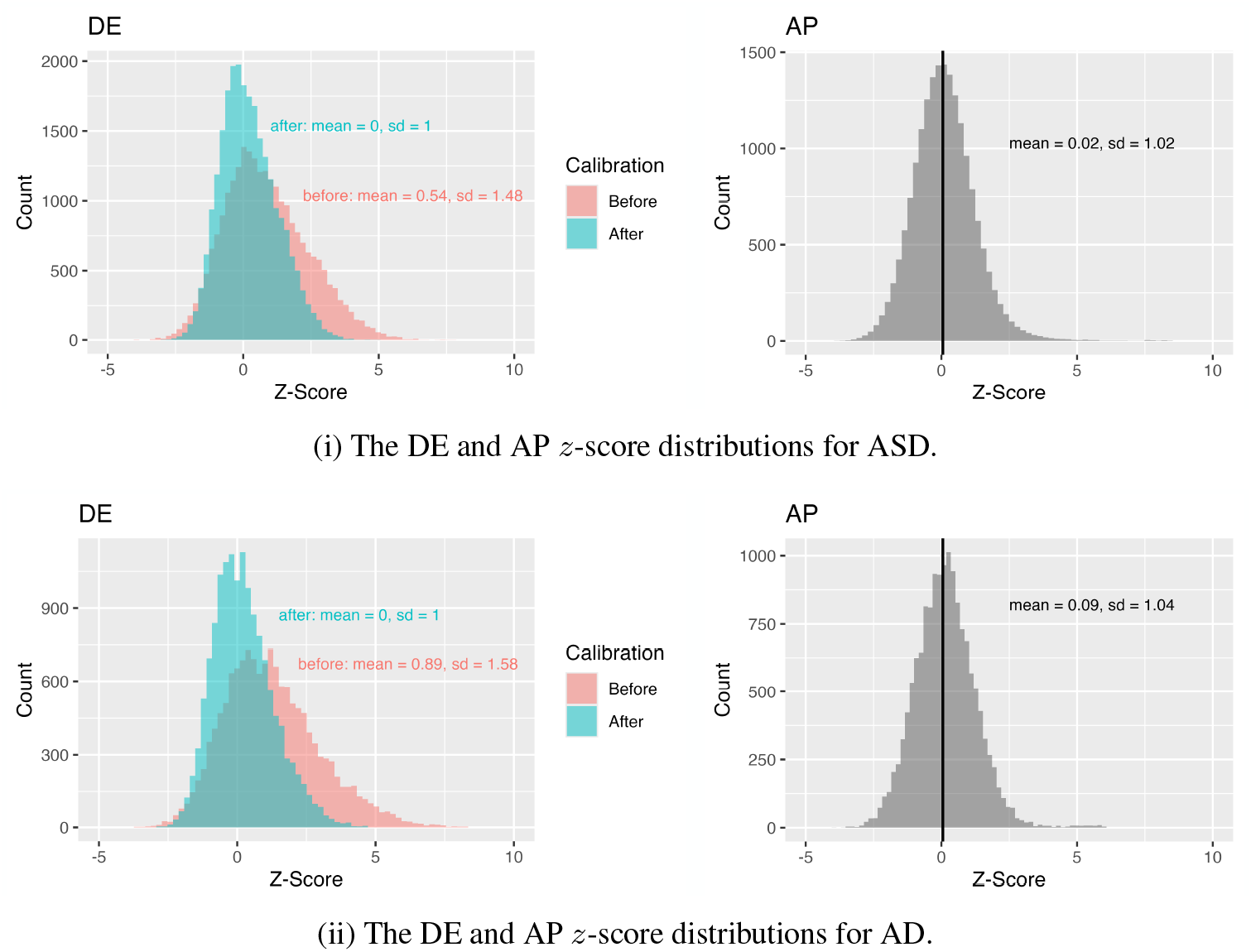
The DE and AP *z*-score distributions for (i) autism spectrum disorder (ASD) and (ii) Alzheimer’s disease (AD). For both ASD and AD, we re-calibrated the DE *z*-scores using Efron’s method [19]. The DE *z*-score distributions before and after calibration can be found in the left panels. The AP *z*-score distributions can be found in the right panels.

After we convert the *p*-values to *z*-scores, we find that some of our *z*-scores are infinite due to the original *p*-values being 0 or very close to 0; for example, we have at least 52 infinite AP *z*-scores for AD from the 52 fixed genes whose *p*-values we set to 0.0. Note: We only have positive infinite values. We replace these infinite *z*-scores by mapping them to equally-spaced values approximately in between the two largest finite *z*-scores in the set. The lower bound of this sequence is determined by taking the second largest finite *z*-score, *a*, and rounding down to the nearest 0.5 value, *a*_floor_ = `floor(`*a*`/2)/2`, and then adding a small amount of noise, *ϵ*_*a*_ = 𝒩 (0, 0.001). The upper bound of this sequence is obtained similarly, by taking the largest finite *z*-score, *b*, and rounding up to the nearest 0.5, *b*_ceiling_ = `ceiling(`*b*`/2)/2`, and then adding a small amount of noise, *ϵ*_*b*_ = 𝒩 (0, 0.001). We then obtain our replacement finite *z*-scores by creating a sequence of equally spaced points between *a*_floor_ and *b*_ceiling_, `seq(from=`*a*_floor_, to=*b*_ceiling_, length.out=numinf+2) where `numinf` is the number of infinite *z*-scores we have to replace, and randomly assigning them to replace the infinite values.

Now that we’ve dealt with our infinite *z*-scores, we can proceed with assessing whether our *z*-scores are well-calibrated and, if they are not, recalibrating them. Using Efron’s `locfdr` method [19], we model and visualize the *z*-score distributions as a mixture of two underlying components: the null (grey) and non-null or signal (purple) distributions. As we can see in Supplementary Figure S1, the AP *z*-scores are distributed well as a mixture of a distribution pretty close to *N* (0, 1) and *N* (*δ, τ*) for both ASD and AD (i.e. well-calibrated). On the other hand, the DE *z*-scores for both appear to have a shifted null distribution, with the maximum likelihood (MLE) estimate for the null distribution being *N* (0.537, 1.477) and *N* (0.892, 1.58) for ASD and AD respectively. Therefore, we recalibrate the DE *z*-scores using the MLE estimate’s from Efron’s method [19]: estimating the mean *µ* and variance *σ* of the distribution of corresponding *z*-values and then adjusted it as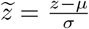. We use these calibrated DE *z*-scores alongside our AP *z*-scores as our input *z*-scores to the joint-HMRF model. Supplementary Figure S2 contains the distributions, means, and standard deviations for all *z*-scores: the DE *z*-scores (before and after calibration) and the unadjusted AP *z*-scores for both ASD and AD.

#### S1.3.2 Gene Filtering and Network Construction

To ensure computational tractability and to focus the analysis on the most biologically relevant genes, the full gene set was reduced before network construction. The first reduction happens by taking the intersection of genes in our three data sources: gene expression data, DE *z*-scores, and AP *z*-scores. Next, we reduce the gene set to those with at least 50% non-zero cells. Prior to network construction via PNS, the gene list was filtered to include only the 8 000 genes exhibiting the highest mean expression level. This selection ensures the retention of robustly expressed transcripts while excluding those with minimal average expression. Gene counts after each of these steps can be found in Supplementary Table S1.

**Table S1:** This table is grouped into three sections detailing: 1) The number of genes after initial filtration and the size of the gene sets within the PNS procedure; 2) The regularization parameter (*λ*) used in PNS; and 3) The size of the constructed networks.

| # | Description | ASD | AD |
| --- | --- | --- | --- |
| **Gene Filtration and PNS Gene Set Sizes** |  |  |  |
| 1 | Data Source Gene Intersection | 15 633 | 11 772<sup>1</sup> |
| 2 | >50% Non-Zero Cells | 13 522 | 11 675 |
| 3 | 8k Genes with Highest Mean Expression | 8 000 | 8 000 |
| 4 | Key Gene Set ($\vert S'\vert $) | 1 532 | 1 518 |
| 5 | Core Gene Set ($\vert S\vert $) | 1 503 | 1 381 |
| 6 | Neighbor Gene Set ($\vert N\vert $) | 5 999 | 5 946 |
| 7 | Final Gene Set ($\vert V\vert $) | 7 502 | 7 327 |
| **PNS Network Construction Parameter** |  |  |  |
| 8 | Regularization Parameter ($\lambda$) | 0.22 | 0.16 |
| **Network Sizes** |  |  |  |
| 9 | PNS Network Size | 13 54 | 2 211 |
| 10 | Pruned Network Size | 1 218 | 2 113 |

<sup>1</sup> This value is slightly inflated due to adding the fixed genes. See Supplementary Section S1.3.1 for more details. The number of intersecting genes for AD without adding the fixed genes is 11 761.

To construct the network we use partial neighborhood selection (PNS); the specifics are laid out in Supplementary Section S1.1.1 and a quick summary with specific thresholds is provided here: The method begins by defining a core gene set *S* by thresholding based on *p*-value and correlation. A starting key gene set is defined by taking the set of genes with *p*-values less than a threshold. Because we have two sets of *p*-values, DE *and* AP, we define the key gene set *S*^*′*^ as the union of genes with a DE or AP *p*-value in the smallest 10% of their respective distributions. To define the core gene set *S*, key genes were subjected to a correlation connectivity filter, retaining only those genes with an absolute correlation of at least *τ* = 0.5. The neighbor gene set *N* is defined by taking the genes in *S*^*C*^ that have an absolute correlation of at least *τ* = 0.5 with at least one gene in *S*. The final gene set is defined by taking the union of the core gene set and the neighbor gene set. The sizes of these gene sets can be found in Supplementary Table S1.

In order to run PNS, we need to set a regularization parameter. We choose this parameter based on a series of metrics including: average degree, *R*^2^, proportion of nodes with degree one, average shortest path, proportion of stable edges, cross-validated mean squared error, and visual structure. A brief description of each metric and its calculation follows:

- Average degree: The average number of edges each node has. We want a well-connected network which corresponds to a larger average degree.
- The coefficient of determination (*R*^2^) or square of the Pearson correlation coefficient for a log–log linear fit of degree frequency vs degree, i.e. how well the network’s degree distribution matches a power-law.
- Proportion of nodes with degree one: The proportion of genes that have degree one. We want a well-connected network which corresponds to a smaller proportion of nodes with degree one.
- Average shortest path: The average shortest path between any two nodes in the network. Calculated using `igraph`’s `mean_distance` function. We want a well-connected network which corresponds to a smaller average shortest path distance.
- Proportion of stable edges: Randomly sample 80% of the genes and use PNS to construct a network. Repeat 100 times and calculate the proportion of edges which appear in at least 60% of the networks. We would like to see a larger proportion of stable edges as it implies a strong underlying signal and structure.
- Cross-validated mean squared error: Obtain the mean squared error from the 5-fold cross-validation on LASSO. A smaller mean squared error is preferred as it implies better fit.
- Visual structure: A qualitative measure based on studying the plotted network. Influenced by network layout. We would like to see a visually rich structure, or a well-connected network (i.e. not sparse) with visually apparent components.

We begin by running PNS and calculating the above metrics on a series of *λ* values with a step size of 0.05: c(0.1, 0.15, …, 0.55, 0.6) – we call this round 1. Based on these metrics, we then hone in on a smaller region to focus on with a step size of 0.01 – we call this round 2. Again, we calculate the above metrics and, based on that, we select a LASSO penalty parameter to construct our final network. For ASD we chose *λ* = 0.22 and for AD we chose *λ* = 0.16, both of which perform reasonably well on the metrics above, including having a visually rich structure, in comparison to other values of *λ*. After obtaining the estimated network following the above processes, we prune the network by removing disconnected components of size 10 or less. This step is performed to minimize network noise and ensure that the analysis focuses only on substantial components, as these smaller subnetworks are often statistically spurious and contain too few genes to meaningfully influence the global inference of the joint-HMRF model.

**Figure S3:**
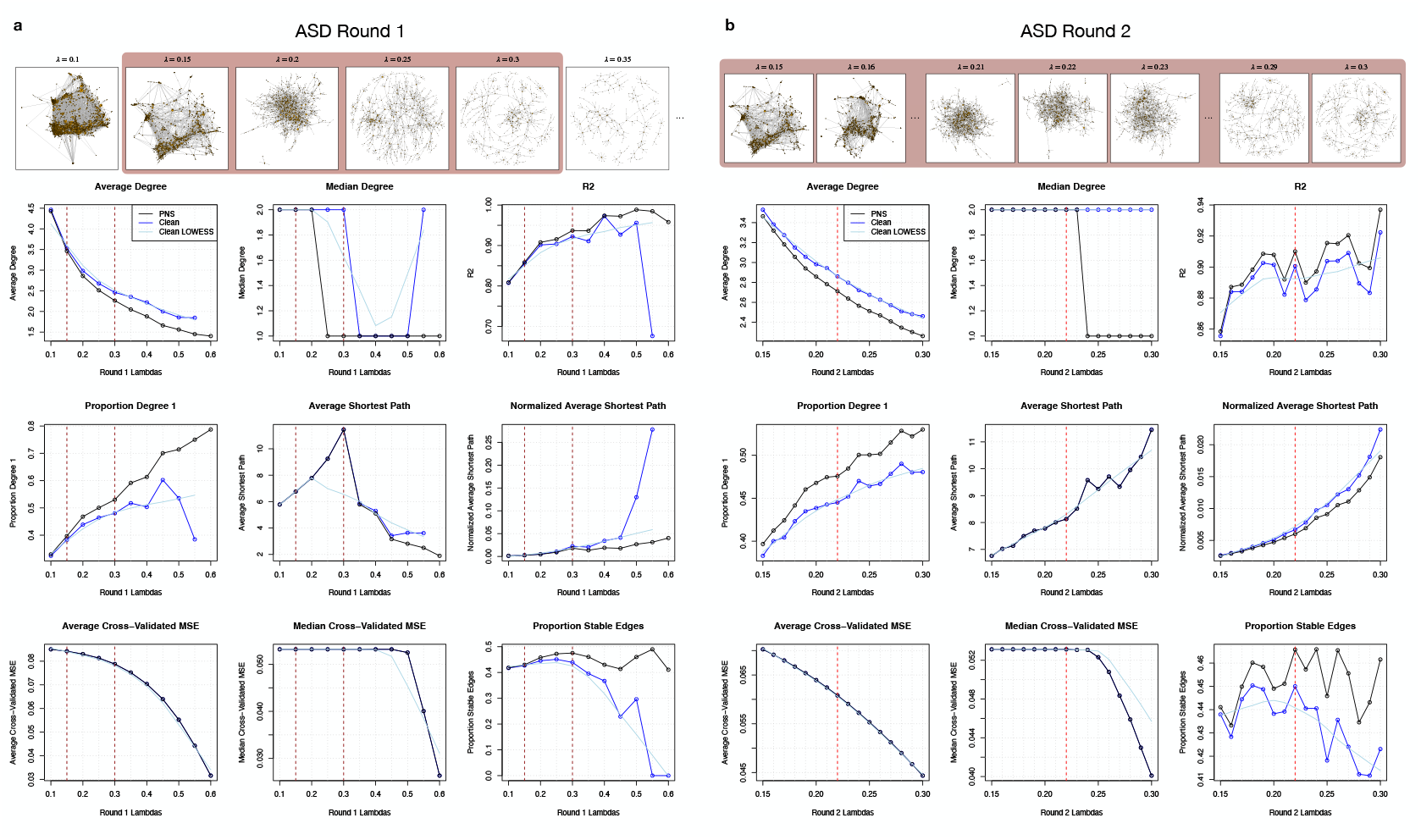
Rounds 1 and 2 of network metrics for *λ* selection for autism spectrum disorder (ASD).

**Figure S4:**
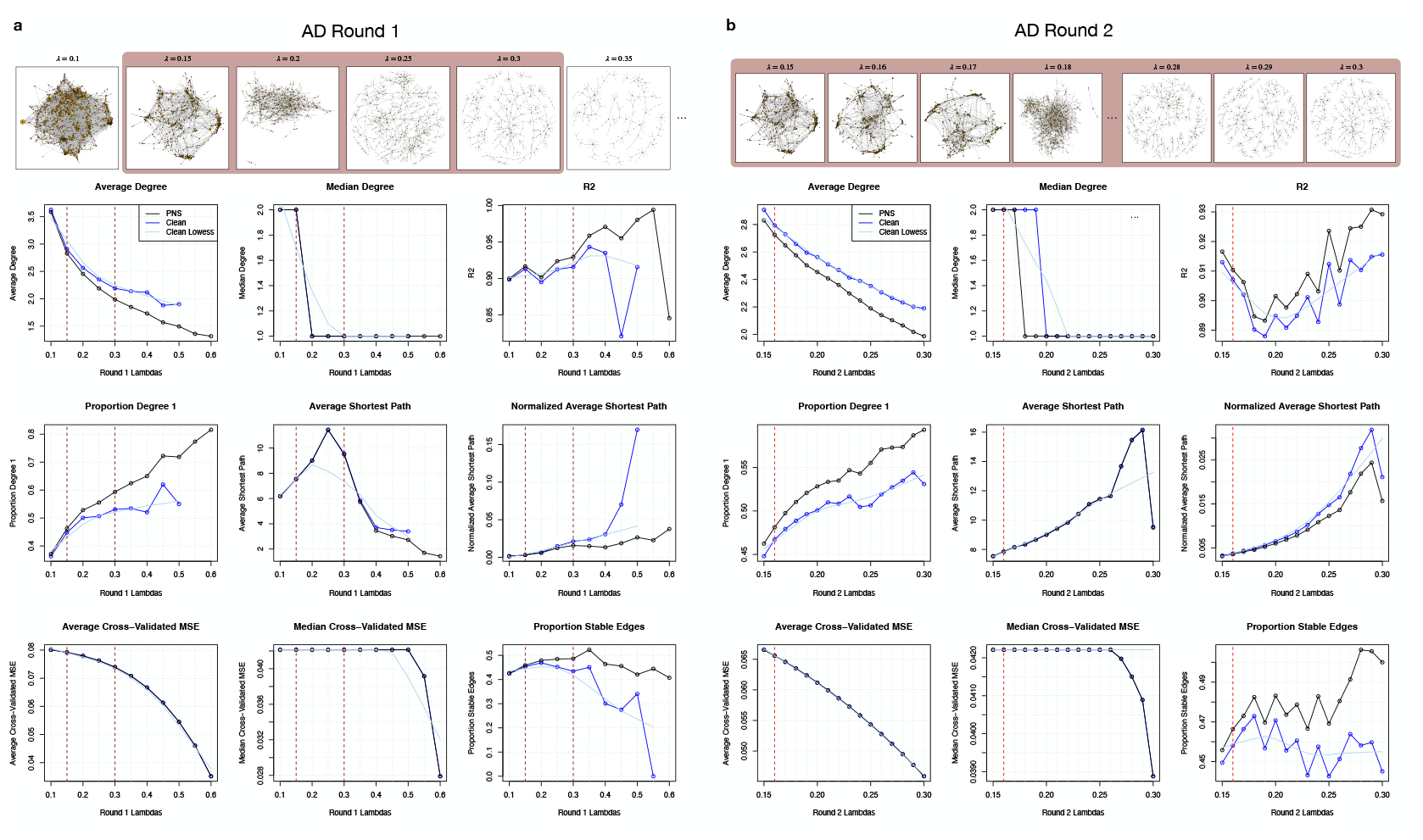
Rounds 1 and 2 of network metrics for *λ* selection for Alzheimer’s disease (AD).

#### S1.3.3 Running Joint-HMRF

As part of the Joint-HMRF algorithm, we begin with two initialization thresholds – one for the DE scores, *C*_*DE*_, and one for the AP scores, *C*_*AP*_ – which determine the genes’ starting states and the normal parameters’ initialization values. Crucially, the initialization must start with a sufficient number of state 1 or active genes at initialization to enable the model to successfully infer any genes as having a hidden state of 1 or active. In particular, this means that there must be a sufficient number of genes with both a DE *z*-score *>* the DE initialization threshold, *C*_*DE*_, and an AP *z*-score *>* the AP initialization threshold, *C*_*AP*_ (Equation 7). However, a range of reasonable values exists for each threshold, *C*_*DE*_ and *C*_*AP*_, that can serve as the initialization. Because the objective surface is non-convex, different initializations may converge to distinct, and sometimes substantially different, results. To ensure stability and robustness of the results, we therefore run the joint-HMRF optimization across a *d*_*DE*_ *× d*_*AP*_ grid of plausible threshold values for 100 iterations. For ASD, our grid was the {0.01, 0.02, …, 0.15}*×* {0.01, 0.02, …, 0.15} grid (symmetric with step-size = 0.01). Because there is less overlapping signal among the AD DE and AP *z*-scores, we have to use a substantially wider and more forgiving range of initialization thresholds: {0.05, 0.10, …, 0.50} *×* {0.05, 0.10, …, 0.50} (symmetric with step-size = 0.05).

From the resulting *d*_*DE*_ · *d*_*AP*_ runs, we then select five runs in a relatively stable region that also contain sufficient active genes. From each of these five runs, we have a set of state labels meaning that, for each gene, we have five possible states. To obtain a gene’s final state label, we take the majority vote amongst the five possible states. If there is a tie we opt to take the lower-valued state, e.g. if there is a tie between state=1 and state=2, we take the gene’s MLHS to be 1.

#### S1.3.4 Stable Two-Step Leiden Clustering

A key component of our clustering method is Two-Step Leiden which is explained in detail in Section S1.1.3. As with many clustering methods, the hyperparameters must be prespecified for both Leiden and Two-Step Leiden. Additionally, Leiden clustering involves randomness in both the initial partition and in the refinement phase of the algorithm, meaning such randomness is also present in Two-Step Leiden. In order to both 1) select clustering hyperparameters resolution *r* and number of clusters *k* and 2) account for randomness in Two-Step Leiden, we add a hyperparameter grid search and run multiple clusterings across random seeds. More specifically, our process is as follows: We start by selecting a set of resolutions and a set of “number of clusters” we want to search over. For each (*r, k*)-pair, we run Two-Step Leiden *s* = 100 times across different random seeds. We then calculate three metrics for each hyperparameter pair across the *s* runs:

- Modularity: Measures the quality of the clustering by comparing the density of edges within communities to the expected density in a randomly rewired network [46]. Calculated for each run separately.
- Silhouette score: Evaluates the quality of the clusters by assessing how similar a data point is to its cluster versus other clusters [47]. This metric requires a dataset for similarity calculation, usually the dataset that the data points were originally clustered using. This is a slightly different setting than we have as our clustering was on a network that was itself constructed from a dataset. We use this original bulk gene expression dataset as the dataset on which similarity is calculated for each data point. This gives a sense of how much these network clusters make sense in the original bulk gene expression data space. Additionally, the silhouette score is calculated per data point. To get a metric for a clustering, we take the average over the silhouette scores of all the data points, giving us a single silhouette score for a set of clusters. Calculated for each run separately.
- Adjusted Rand Index (ARI): Measures the similarity between pairs of clusterings [48]. We calculate the ARI for each pair of runs and take the mean. This aggregated value gives us a sense of the stability of the runs for a pair of hyperparameters. Calculated for each pair of runs and then averaged.

We then combine the three metrics into an aggregated score to select the “best” (*r, k*)-pair. To ensure equal weight, we scale all the metrics to be between 0 and 1 by type, e.g. scale all the modularity scores to be between 0 and 1 where the smallest modularity is now 0 and the largest is 1. The scaled metrics are added together to obtain the final aggregated clustering score which factors in modularity, how well the clusters fit the data, and stability. The best hyperparameter pair, (*r*^*′*^, *k*^*′*^), is picked by maximizing this score.

Final clusters were obtained using consensus clustering. For the selected hyperparameters, we took the *s* runs and constructed a consensus matrix *M* where *M*_*ij*_ is the proportion of times genes *i* and *j* co-occurred in the same cluster. We then applied hierarchical clustering to *M* to obtain the final *k*^*′*^ clusters.

This method of clustering does leave us with a couple large clusters for both applications. Thus, large clusters (defined as clusters with 250 or more genes) were subsequently examined and manually subdivided to capture finer-grained structure. For ASD, we simply cut each large cluster into two or three components based on the structure of the dendrogram. For AD, because of the large clusters’ dendrograms’ diffuse structure and lack of distinct cluster boundaries, we used WGCNA to subdivide the large clusters [7].

#### S1.3.5 Cluster Labeling

Next we analyze the composition of the clusters obtained from our clustering procedure. In particular, remember that our goal is to determine which p-reactive genes are likely to actually be active, i.e. an AP gene in addition to being a DE gene. We do this by labeling clusters based on the composition of active versus p-reactive genes. These designations split gene clusters into those more likely to play a fundamental role in the development of the phenotype (etiological) versus those more likely just differentially expressed as a consequence of the phenotype (emergent).

For each gene state and cluster there are two possible views: the proportion 1) within a cluster or 2) across that state. For example, say a cluster has *b* genes total and *a* of a certain gene state and that there are *c* total genes of that state across all clusters. Then we consider both the proportion of that gene state within this cluster, 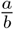, and the proportion of that gene state within this cluster compared to the total number of genes labeled as this state, 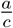. We define a set of application-specific cascading if-else rules based on both 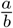 and 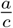 thresholds. These rules are defined by studying these proportional values, placing more emphasis on the values related to the active genes followed by those related to the p-reactive genes. Our sets of rules for ASD and AD can be found in Supplementary Table S2, where the sub-column ***State*** thresholds correspond to the 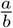 value and the sub-column ***Cluster*** thresholds correspond to the 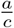 value. A cluster is designated with the label if *either* its state proportion (***State*** or 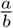) *or* its cluster proportion (***Cluster*** or 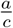) exceeds the corresponding threshold listed in the cell. For example, if a cluster we’ve identified in the ASD application has either over 15% of all the active genes in that cluster or is comprised of over 20% active genes, it is designated as etiological.

**Table S2:**
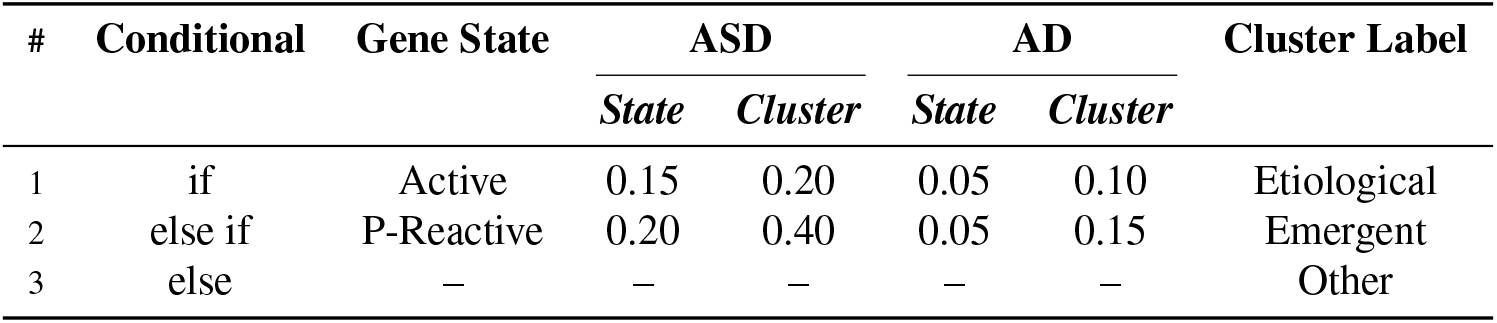
If-else rules for ASD and AD for labeling clusters as Etiological, Emergent, and Other. The rules proceed in a waterfall format, applied in order from #1 to #3. A cluster is designated with the label if *either* its state proportion (***State***) *or* its cluster proportion (***Cluster***) of the corresponding state exceeds (*>*) the corresponding threshold value listed in the cell.

| # | Conditional | Gene State | ASD |  | AD |  | Cluster Label |
| --- | --- | --- | --- | --- | --- | --- | --- |
|  |  |  | *State* | *Cluster* | *State* | *Cluster* |  |
| 1 | if | Active | 0.15 | 0.20 | 0.05 | 0.10 | Etiological |
| 2 | else if | P-Reactive | 0.20 | 0.40 | 0.05 | 0.15 | Emergent |
| 3 | else | — | — | — | — | — | Other |

### S2 Supplementary Results

#### S2.1 Cell Type-Specific Marker Gene Enrichment

To assign biological context to our clusters, we leveraged cell type-specific marker genes derived from single-nucleus RNA-sequencing (snRNA-seq) data [20] to establish which clusters are enriched for which cell type’s marker genes. The Velmeshev et al. [20] dataset comprises 413 682 nuclei across 108 cortical tissue samples from 60 prenatal and postnatal human individuals with no neuropathological abnormalities. 358 663 nuclei were retained after removing a cluster of cell debris. Additionally, the authors integrated their data with published datasets of prenatal and postnatal human cortical development. To match the age range of our expression profiles for gene network construction, we subsetted this snRNA-seq dataset to include only cells derived from individuals aged 2 years and older. Within our snRNA dataset, we are provided the following cell types: interneurons, excitatory neurons, microglia, pericytes, vascular cells, fibrous astrocytes, protoplasmic astrocytes, glial progenitors, oligodendrocyte progenitor cells (OPC), and oligodendrocytes. We combined the fibrous and protoplasmic astrocytes into one broad astrocyte cell type. We excluded the glial progenitors from our analysis due to their low cell count (*n* = 87 within this age range) and uncertain lineage (oligodendrocyte vs astrocyte), which prevented a reliable combination with another cell type. The breakdown of cell counts by cell type after subsetting by age and combining/excluding cell types is presented in Supplementary Table S3. As one can see, we do have a wide range of counts per cell type, with the smallest cell type being pericytes (1 284) and the largest cell type being excitatory neurons (58 984)

**Table S3:** Cell Counts by Major Cell Type. This table provides the cell-level breakdown of the single-nuclei RNA-sequencing (snRNA-seq) dataset used for identifying cell-type specific marker genes. The dataset was filtered by age (2 years and older) and sub cell types were combined or excluded (combined: astrocytes, excluded: glial progenitors). Counts are partitioned across the eight major cell types identified. Abbreviations: IntNeu, Inhibitory Neurons; ExcNeu, Excitatory Neurons; Ast, Astrocytes; OPC, Oligodendrocyte Precursor Cells; Oligo, Oligodendrocytes; Vasc, Vascular Cells.

| IntNeu | ExcNeu | Microglia | Ast | OPC | Oligo | Pericytes | Vasc | Total |
| --- | --- | --- | --- | --- | --- | --- | --- | --- |
| 29 951 | 58 984 | 9 937 | 24 551 | 17 138 | 40 314 | 1 284 | 2 323 | 184 482 |

We log-normalize our data by dividing each gene by its library size, then dividing by 10 000 to get the normalized counts per ten thousand (CP10K), and then taking the base 2 logarithm with a pseudocount of 1: log_2_(CP10K + 1). To identify marker genes, we use `Seurat`’s `FindMarkers` [21, 22] using the Wilcoxon Rank Sum test (`test.use=‘wilcox’`) for computational reasons and the lack of distributional assumptions. We also set the following arguments: `only.pos=TRUE`, `logfc.threshold=0`, `min.pct=0`, `max.cells.per.ident=5 000` in order to identify all possible positive marker genes and to deal with the imbalance in cells across cell types (the other arguments were left as the default). Because we set `max.cells.per.ident=5 000`, when an identity (cell type or all cell types but one) has more than 5 000 cells, `FindMarkers`’ downsamples the cells to 5 000. This sampling is random (set by a random seed) and results can change greatly across different subsamples. Thus, to ensure stability, we aggregate marker genes across 10 runs with unique random seeds for each run and cell type combination. For run *i*, we run `FindMarkers` on each cell type, *c*, with a unique random seed (`random.seed =` *s*_*ic*_). After obtaining the potential marker genes, we filter the potential markers by dropping all genes with an adjusted *p*-value ≥0.05. The marker-finding functions in `Seurat` do not impose exclusivity; that is, they do not restrict marker genes to be unique to a single cell type. To enforce this standard uniqueness criterion for marker genes, if a gene was identified in multiple cell types, we only keep the instance with the largest logFC. Finally, we further reduced the potential marker gene set by dropping any gene with a logFC ≤1. We repeat this procedure ten times, resulting in ten sets of cell type-specific marker genes. For our final, stable marker gene set, for each cell type, we take the set of marker genes which appear in *>* 5 runs.

After identifying marker genes for seven major cell types (interneurons, excitatory neurons, microglia, astrocytes, oligodendrocytes, pericytes, and vascular cells), we tested each of the clusters for cell type-specific marker gene enrichment for both applications (Figures 3d, 5d). The marker genes can be found in Supplementary Tables S6 and S13 and the enrichment results can be found in Supplementary Tables S7 and S14.

### S2.2 Comparison with Gandal et al. [1] WGCNA modules

We also compare our identified clusters with the 35 WGCNA modules identified by Gandal et al. [1]. They assigned cell-type identities to the WGCNA modules using the Expression Weighted Cell Type Enrichment (EWCE) method [23], which assesses the enrichment of gene sets by evaluating whether a set of genes has a higher expression within a cell type than expected by chance, alongside a scRNA-seq reference dataset from Lake et al. [49]. The majority of their WGCNA modules are only significantly associated with one cell type, however, seven of the modules are associated with two cell types (M3, M5, M26, M22, M7, M10, M24). Figures S5 and S6 show the Jaccard similarity index between our identified clusters (for ASD and AD) and the WGCNA modules. More specifically, cell_*ij*_ corresponds to the Jaccard index of cluster *i* and module 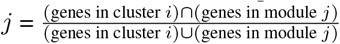.

**Figure S5:**
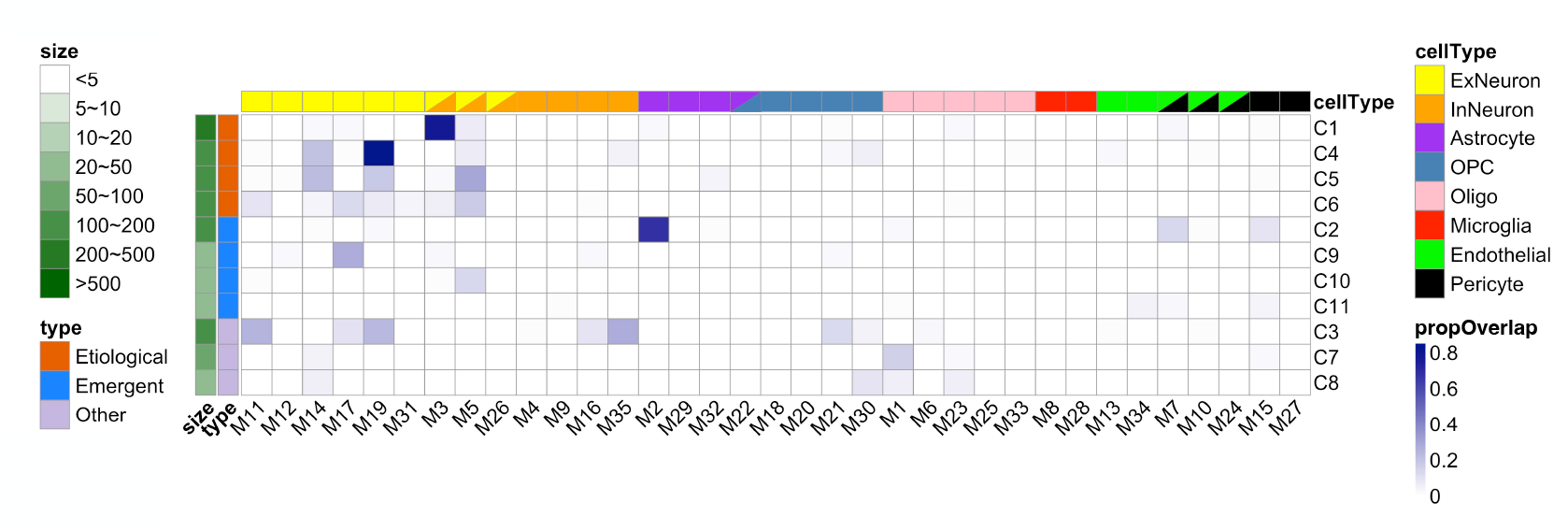
The comparison between our autism spectrum disorder (ASD) DAWN-SCAPE clusters and the WGCNA modules found in [1]. The heatmap displays the Jaccard index between the DAWN-SCAPE clusters and the WGCNA modules.

For ASD, we can see that our identified etiological clusters have the highest similarity with the neuron-type (e.g. excitatory neurons and and interneurons) WGCNA modules, matching our cell type-specific marker gene enrichment analysis results with an external single-nucleus dataset in Figure 3d. Additionally, while the emergent clusters have some similarity with the neuron-type WGCNA modules, they are also similar to some non-neuron type (e.g. astrocytes, endothelial, pericytes) WGCNA modules. It is found in previous studies that ASD genes are more enriched in neuron-type of cells [13], which concurs with our results.

**Figure S6:**
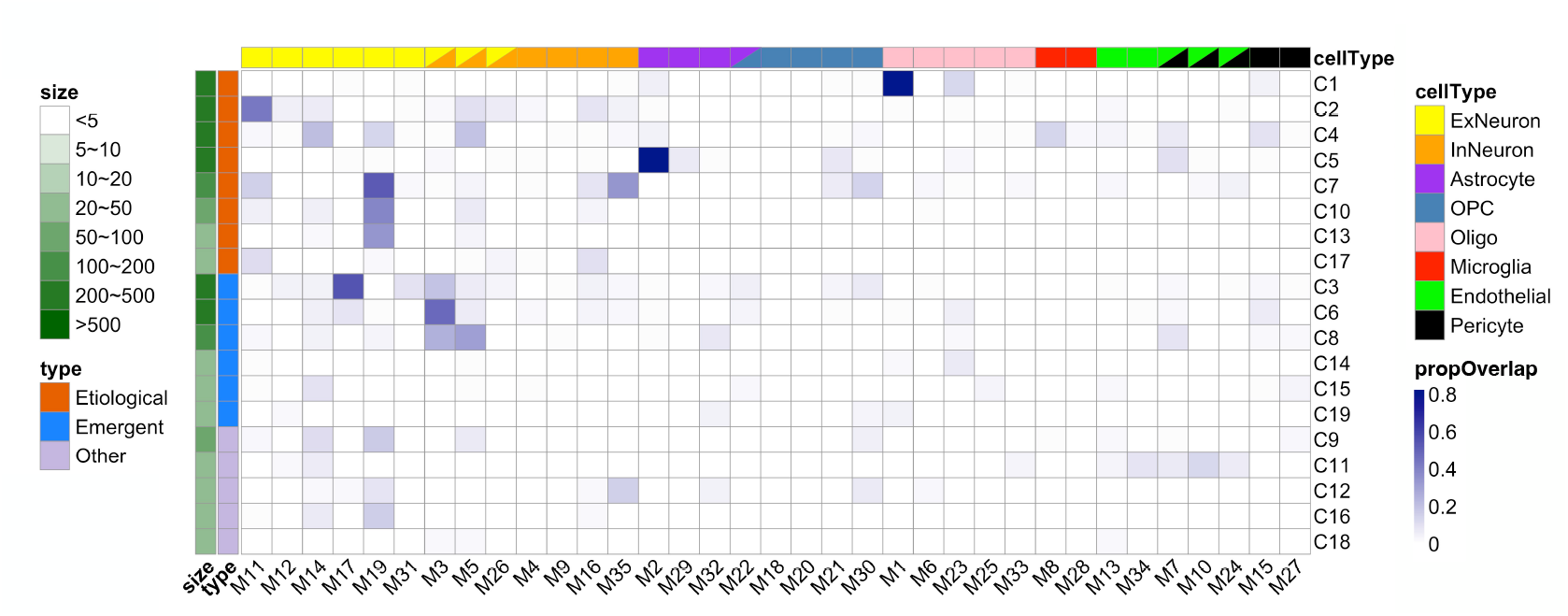
The comparison between our Alzheimer’s disease (AD) DAWN-SCAPE clusters and the WGCNA modules found in [1]. The heatmap displays the Jaccard index between the DAWN-SCAPE clusters and the WGCNA modules.

For AD, we see that while our etiological clusters also have high similarity with the neuronal WGCNA modules, there is also high similarity between our astrocyte etiological cluster (C5) with a WGCNA astrocyte module (M2) and between our oligodendrocyte etiological cluster (C1) with a WGCNA oligodendrocyte module (M1) Figure 5d. The emergent clusters also have the most similarity with the neuronal WGCNA modules and we don’t see a strong similarity with any of the other WGCNA cell type modules. Overall we can see that the Gandal et al. [1] WGCNA modules and our DAWN-SCAPE clusters do roughly align on the broad cell type associations with a couple of cluster *×* module pairs of the same cell types having extremely high similarities. This provides additional support for the robustness of our cell type-specific associations and the validity of our PNS clustering approach.

**Figure S7:**
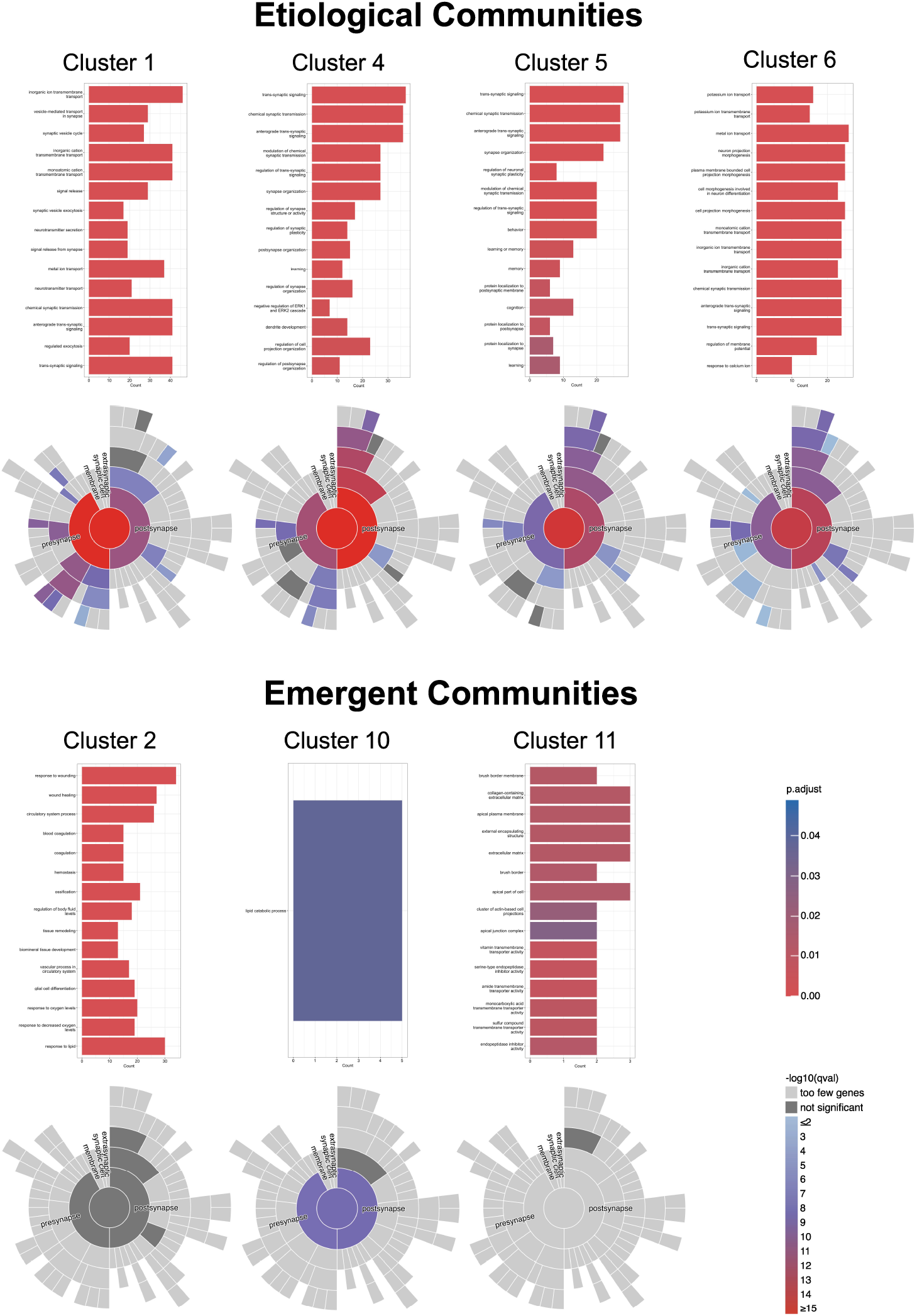
Gene Ontology (GO) enrichment results for autism spectrum disorder (ASD) etiological and emergent clusters. Only the top 15 enrichment terms by adjusted *p*-value are displayed. See Supplementary Table S10 for the full set of GO enrichment terms.

**Figure S8:**
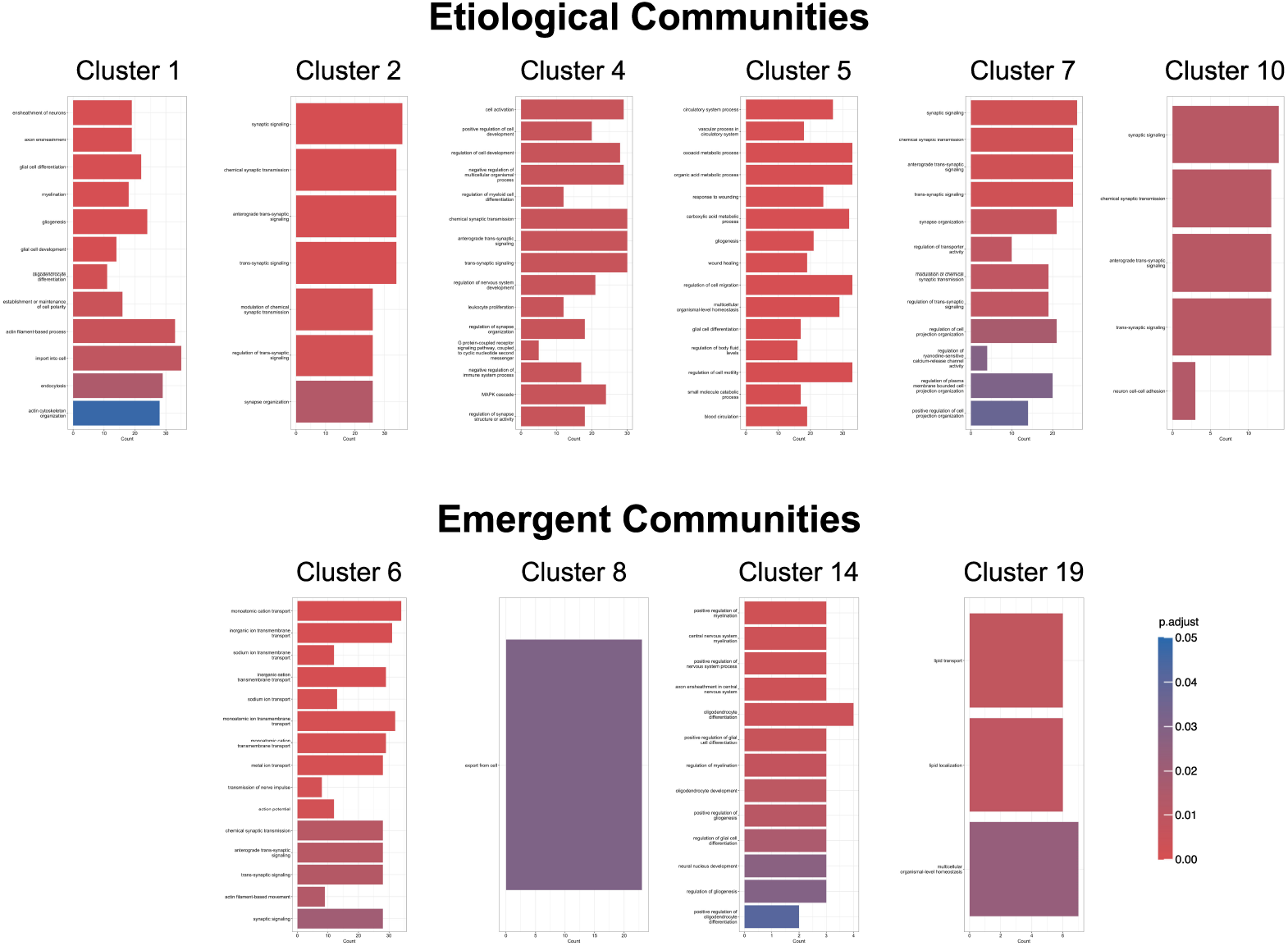
Gene Ontology (GO) enrichment results for Alzheimer’s disease (AD) etiological and emergent clusters. Only the top 15 enrichment terms by adjusted *p*-value are displayed. See Supplementary Table S17 for the full set of GO enrichment terms.

**Figure S9:**
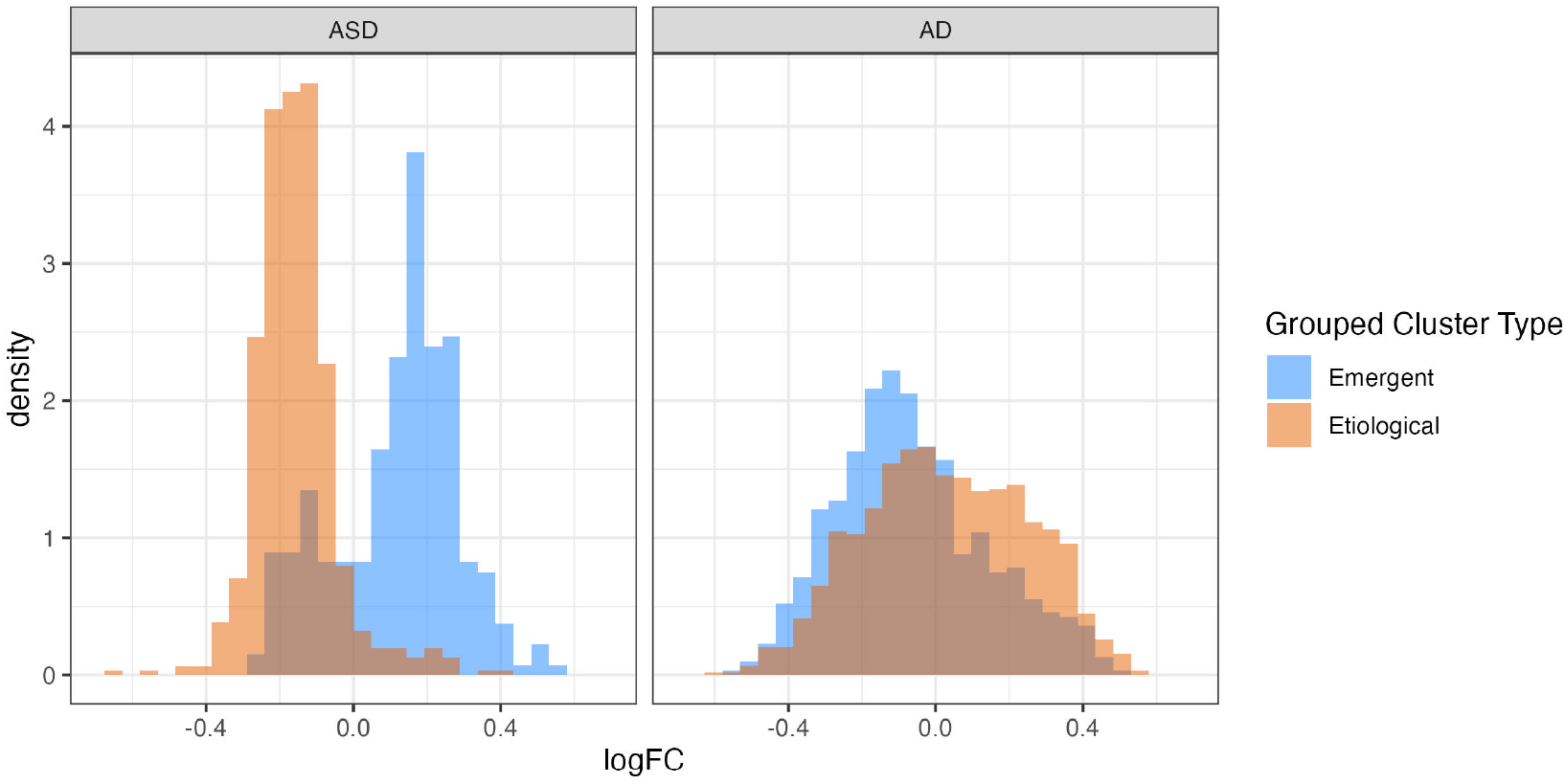
The whole-cortex differential expression of genes in etiological vs emergent clusters for autism spectrum disorder (left) and Alzheimer’s disease (right). The log fold change (logFC) for each gene is shown. Positive logFC means a gene is more expressed while negative means less expressed.

**Figure S10:**
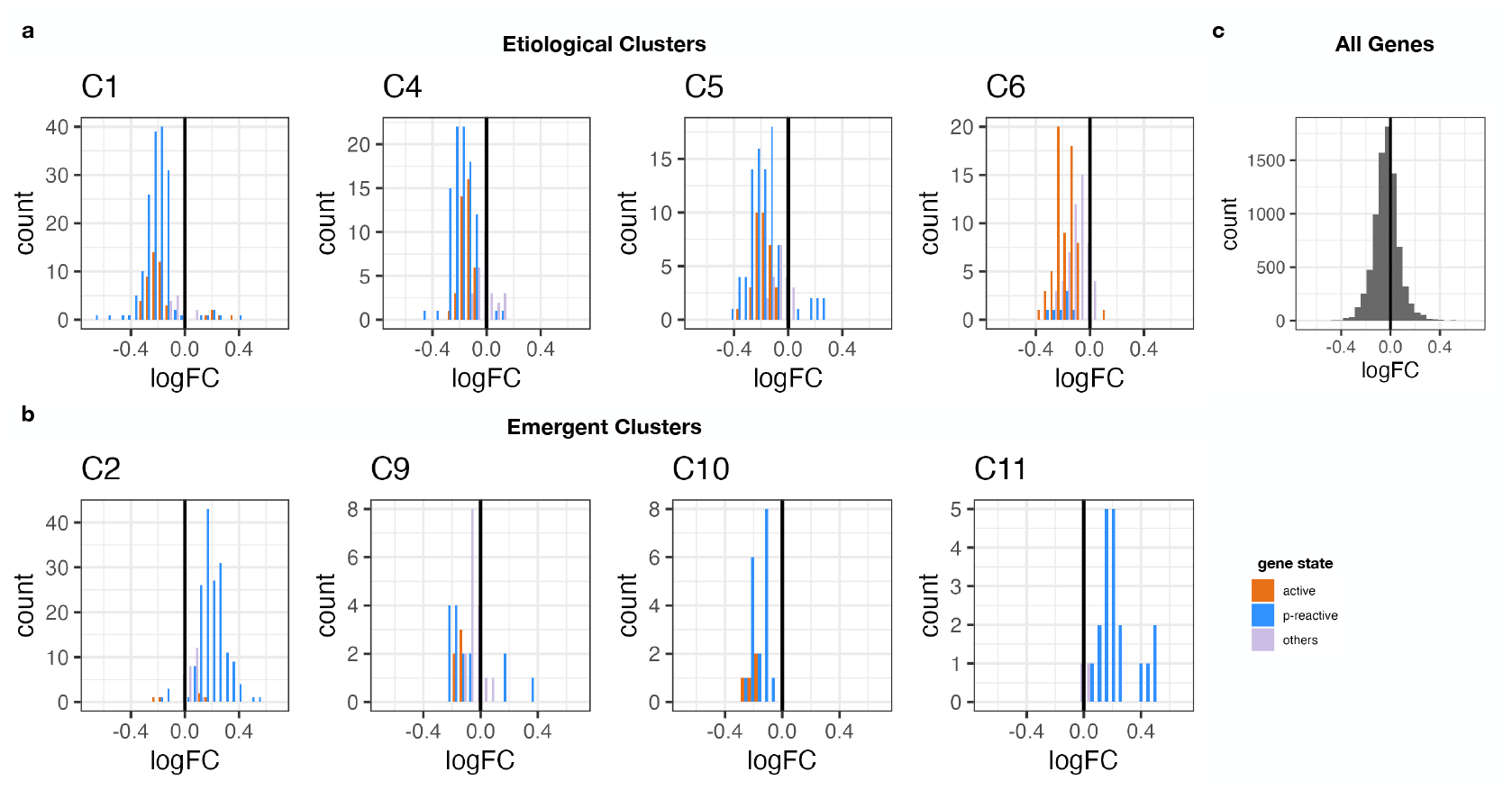
The distribution of the log fold change (logFC) for whole-cortex differentially expressed genes in autism spectrum disorder by cluster. The distributions of genes in etiological clusters (C1, C4, C5, C6) are shown in (a), those of emergent clusters (C2, C9, C10, C11) are shown in (b), and the overall logFC distribution of all 8 000 genes considered for network construction is shown in (c). Positive logFC indicates higher gene expression in AD tissue, while negative logFC indicates lower expression.

**Figure S11:**
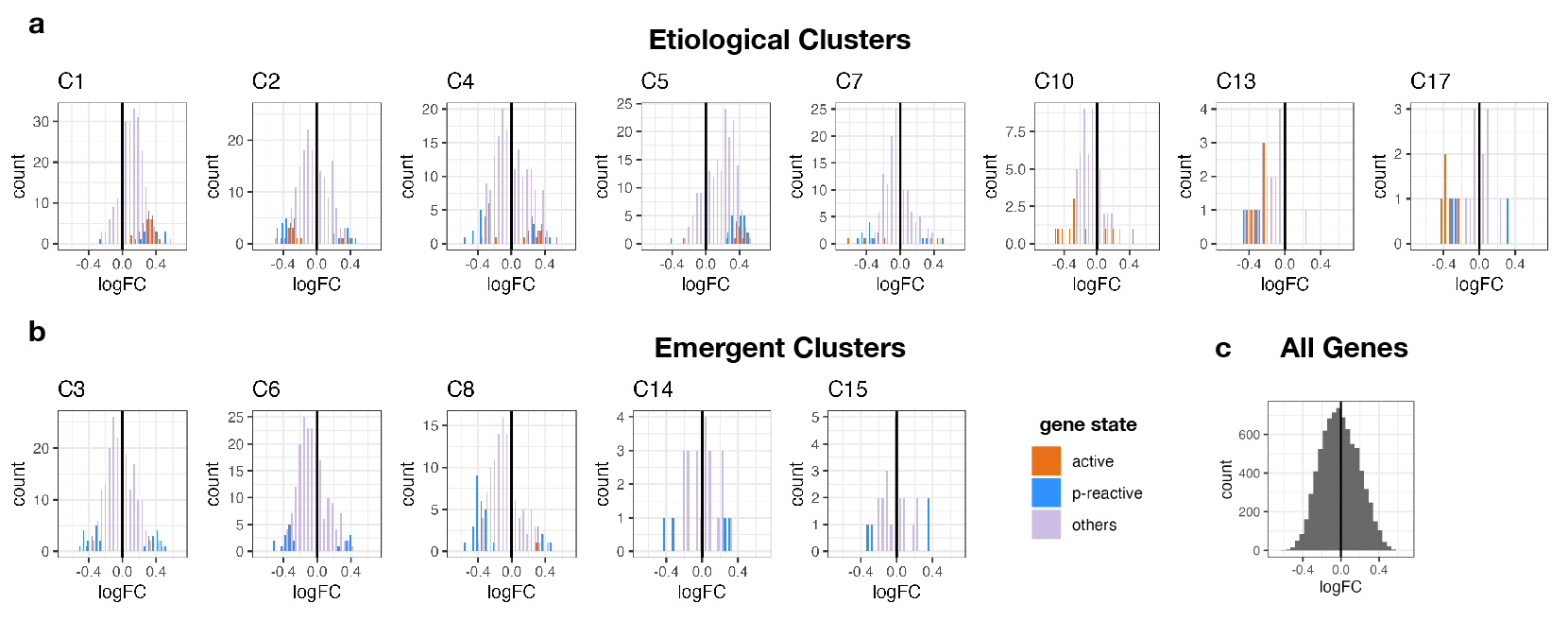
The distribution of the log fold change (logFC) for whole-cortex differentially expressed genes in Alzheimer’s disease (AD) by cluster. The distributions of genes in etiological clusters (C1, C2, C4, C5, C7, C10, C13, C17) are shown in (a), those of emergent clusters (C3, C6, C8, C14, C15) are shown in (b),and the overall logFC distribution of all 8 000 genes considered for network construction is shown in (c). Positive logFC indicates higher gene expression in ASD tissue, while negative logFC indicates lower expression.

## Notes

### Competing Interest Statement

The authors have declared no competing interest.

### Funding

- NIMH Repository and Genomics Resource: MH11165803
- Simons Foundation: SFARI SF1018804

